# Ion selectivity and activation mechanism for kalium channelrhodopsins

**DOI:** 10.1101/2023.07.22.550149

**Authors:** Mingfeng Zhang, Yuanyue Shan, Liping Zhao, Xiao Li, Duanqing Pei

## Abstract

Channelrhodopsins harvest the light and convert photons to the cellular ion flow. The ion selectivity and activation mechanism at the atomic level remains unknown. Here we describe cryo-EM structures for *H. catenoides* kalium channelrhodopsin (*Hc*KCR1), its paralog, sodium selective channelrhodopsin (*Hc*CCR), an open state of *Hc*KCR1 (C110T), the voltage-dependent inwardly rectifier (D116N) and higher potassium selective channelrhodopsin (*B1*ChR2) from *Bilabrum sp*, illuminating the ion selectivity and activation mechanism. Briefly, the hourglass shaped lumen is occupied by the stepwise dehydrated potassium in both intracellular and extracellular side. The aromatic amino acids likely function as partial dehydrated potassium filter in the extracellular lumen, and intracellular dehydrated ion occupying layer chooses the right size of dehydrated ion, thus specifying ion selectivity and the higher dehydration capacity, the higher potassium selectivity. Furthermore, structural comparison of *Hc*KCR1 and C110T suggested that the conformational changes of retinal triggers the extracellular side of TM6 extension as well as the retinal interaction residues motion, which then leads to ion flow. Our results not only uncovered the ion selectivity mechanism of potassium or sodium selective channelrhodopsins, but also elucidated their activation mechanism. It may provide a framework for designing next generation optogenetic tools.

## Main

## Introduction

Channelrhodopsins are light-activated ion channels that have extensively been applied for controlling cell excitability^1^. They belong to the microbial rhodopsin superfamily and the first channelrhodopsin was discovered in green algae, *Chlamydomonas reinhardtii*^2^. After the hallmark discovery, several cation/anion-conducting channelrhodopsins (CCR/ACR) variants have been identified from natural resources^3–7^ or been designed^8–12^, which greatly expand the optogenetics toolbox for cell excitability control.

Excitingly, the channelrhodopsins from *H. catenoides* (*Hc*KCR1 and *Hc*KCR2)^13^ are identified as potassium selective channelrhodopsins, which not only extend our view of potassium selectivity but also can help us to rational control cell excitability. Furthermore, *Wobblia lunata* (*Wi*ChR), *Bilabrum* sp. (*B1*ChR2), and *Colponema vietnamica* (*Cov*KCR1 and *Cov*KCR2)^14^ exhibit higher potassium selective than the *Hc*KCR1 and *Hc*KCR2, which illuminate the bright prospect to engineer as the ultra-high potassium selective channelrhodopsins that can well control neuronal inhibition.

Interestingly, another channelrhodopsin paralog (*Hc*CCR), from in *H. catenoides,* with 70-73% sequencing similarity to *Hc*KCRs, is sodium selective^15^, suggesting that these channelrhodopsins may also be engineered to desired ion selectivity to control cellular specific ions. However, these potassium selective channelrhodopsins lack the canonical conserved sequence “T(S)VGY(F)G”, known as K^+^ selectivity filter^16^. The lack of basic understanding on principles specifying ion selectivity would hamper the design of next generation of channelrhodopsins for optogenetics.

In this study, we show that *Hc*KCR1 is strictly potassium selective like the traditional tetrameric potassium channel and we solved 2.6 Å cryo-EM structures of wildtype *Hc*KCR1 and 2.5 Å cryo-EM structures of open state of *Hc*KCR1 in potassium environment. The high resolution cryo-EM structure of *Hc*KCR1 and its open state indicate that the entire continuous potassium permeation pathway and the conformational changes of retinal as well as the key gating elements. The continuous potassium from both extracellular side and intracellular side of *Hc*KCR1 exhibited the stepwise dehydrating manner, and the dehydrated potassium gradient is safeguarded by the aromatic amino acids in the extracellular side. Strikingly, the 2.4 Å cryo-EM structures of sodium selective channelrhodopsin *Hc*CCR in the sodium environment reveals that the sizes of extracellular hydrated ion occupied lumen and intracellular dehydrated ion occupied lumen select the appropriate sodium to flow. Interestingly, the 2.7 Å cryo-EM structures of higher potassium selective channelrhodopsin *B1*ChR2 in the potassium environment suggest that the higher dehydration capacity, the higher potassium selectivity.

## Results

### *Hc*KCR1 is strictly potassium selective like the traditional tetrameric potassium channel

To gain insight into the interesting high potassium selective channelrhodopsins, we overexpressed C-terminal GFP tagged *Hc*KCR1 in HEK293T cells and performed the whole cell patch clamp under the physiological sodium versus potassium gradient conditions (Fig. 1a). The *Hc*KCR1 transfected cells can generate reproducible and giant photoactivated potassium currents with a reversal potential of-83.8 ± 5.4 mV for peak currents and-75.0 ± 2.8 mV for steady currents (Fig. 1b, 1d and 1f), which are consistent with the previous report^13^. Because the physiological environment produces a native potassium (intracellular) versus sodium (extracellular) gradient, one could explain that the potassium selectivity of *Hc*KCR1 could be due to the direction of ion flow. We used the inside-out patch method to invert the physiological potassium ion gradient (Ion in intracellular side is sodium while the ion in extracellular side is potassium) to test the speculation (Fig. 1a). As *Hc*KCR1 can generate sufficiently high inside out photoactivated microscopy currents, the reversal potential of *Hc*KCR1 can be precisely determined with-90.8 ± 2.9 mV for peak currents and-81.5 ± 2.4 mV for steady currents (Fig. 1c, 1e and 1f) and based on the Goldman-Hodgkin-Katz Equations (GHK)^17^, the P_K_/P_Na_ permeability ratio of *Hc*KCR1 is around 33 for the peak currents and 23 for the steady currents under the inverted physiological potassium gradient, which resembles the physiological sodium versus potassium gradient conditions. Therefore, physiological sodium and potassium gradient conditions are not responsible for potassium selectivity, and *Hc*KCR1 is a strictly potassium selective ion channel, just like the traditional tetrameric potassium channel^16^.

**Fig. 1.**
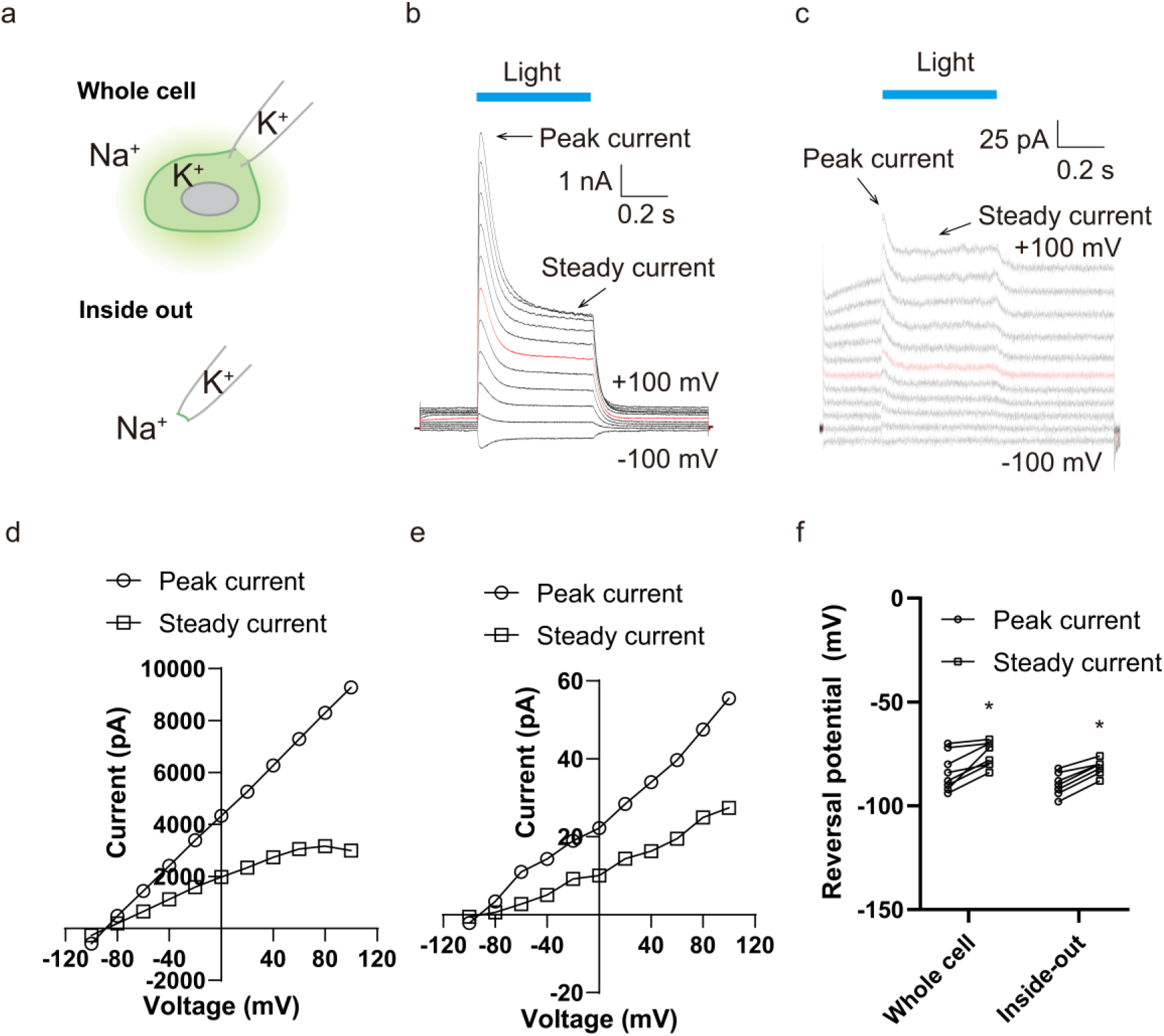
Electrophysiological results of *Hc*KCR1. **a**, Cartoon representation of the whole cell and inside-out mode for recording the light activated currents. **b,** The representative physiological whole cell electrophysiological recordings (out Na and in K) from the HEK293T cell expressing wildtype *Hc*KCR1 with the voltage-pulse from-100 mV to +100 mV with a +20 mV step. Each step is stimulated by blue light and that is indicated by the blue line. The red trace stands for the current recorded at the holding potential of 0 mV. As the channel undergoes the fast inactivation process, the peak currents and inactivated steady currents are indicated. **c,** The recoding protocol is consistent with the whole cell mode, while the currents are recorded at the inverted physiological condition (out K and in Na). **d-e,** The current-voltage (I-V) plot of light activated peak and steady currents of whole cell mode **(d)** from **(b)** and inside out mode **(e)** from **(c)**. **f,** The reversal potential of peak current and steady current of whole cell and inside out mode. The statistic data points are the Mean ± SEM (n is at least 5 cells for each variant). *, p < 0.05 by the student’s t-test

### Determination of HcKCR1 structures through Cryo-EM

Since the *Hc*KCR1 is a strictly potassium selective ion channel, the potassium must be occupied in the ion permeation pathway. To dissect the potassium selectivity mechanism, we purified *Hc*KCR1 in potassium and digitonin environment and then subjected it into the routine cryo-electron microscopy (cryo-EM) process. After sample preparation, image acquisition and data processing, we obtained the near atomic *Hc*KCR1 map at 2.6 Å resolution (Extended Data Fig. 1a-c). The high-resolution map of *Hc*KCR1 allowed us to build molecular models of amino acids as well as the lipids and potential ions/waters (Extended Data Fig. 2).

**Fig. 2.**
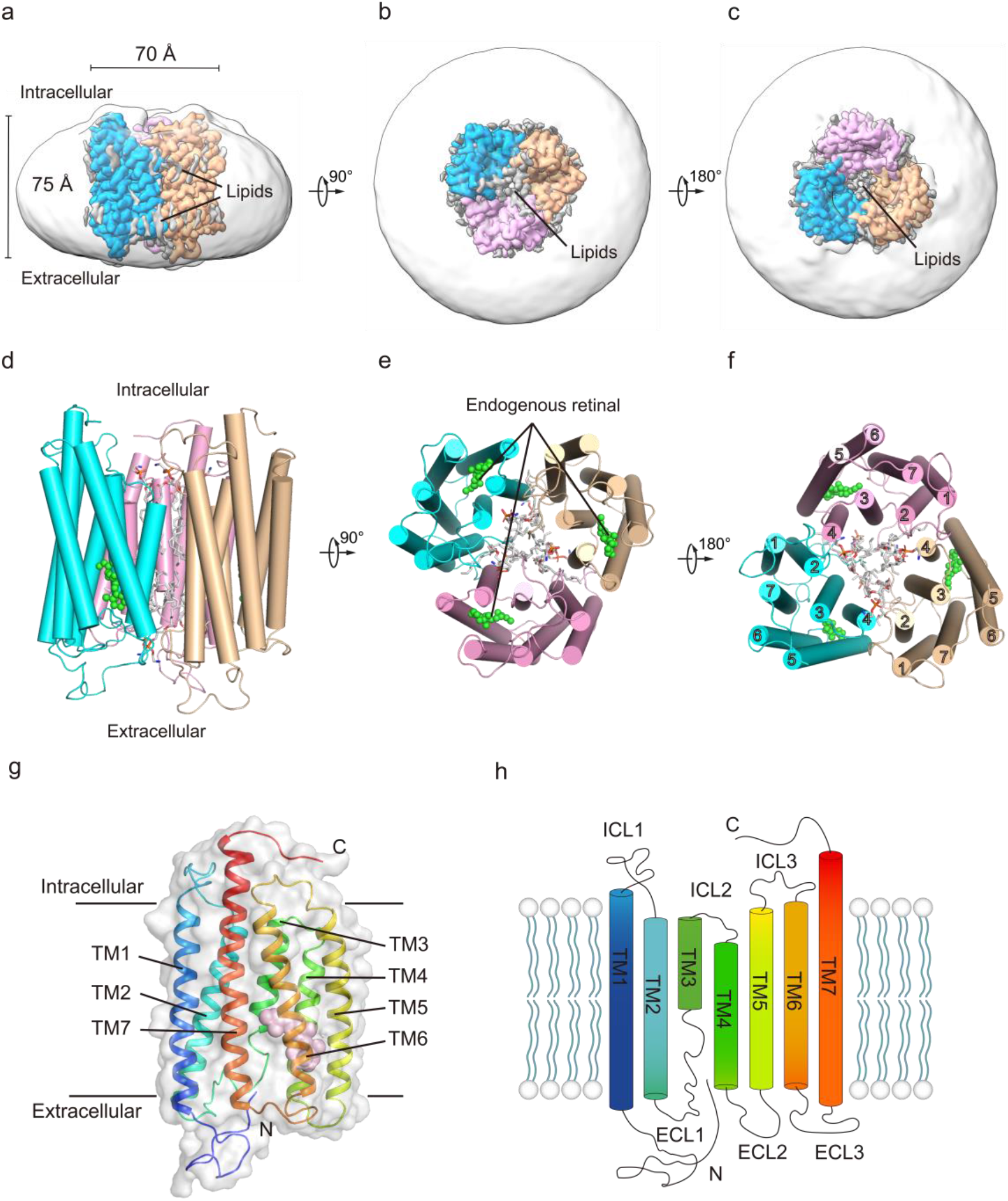
Overall structure of trimeric *Hc*KCR1. **a-c**, Cryo-EM density map of trimeric *Hc*KCR1 bounded with endogenous retinal viewed from the membrane plane (**a**), the extracellular side (**b**), and the intracellular side (**c**). Subunits are colored teal, pink, or wheat; the central pore lipids are colored grey; and the detergent micelle is transparent. **d-e,** Model shown in the same views as **a-c** respectively with endogenous potential all-trans retinal (ATR) shown as green spheres. **g-h,** Model (**g**) and cartoon (**h**) of a single *Hc*KCR1 subunit colored blue (N-terminus) to red (C-terminus).

*Hc*KCR1 adopts trimer architecture like ChRmine^18, 19^, instead of dimer formation of ChR2 or Chrimson^20, 21^ known for light activated non-selectivity cation channels (Fig. 2a-f). Each subunit consists of the canonical rhodopsin topology with seven transmembrane helices (TM1-TM7), three intracellular linkers (ICL1-ICL3), three extracellular linkers (ECL1-ECL3), an extracellular N-terminal domain and an intracellular C-terminal domain (Fig. 2g-h). Each monomer is connected by amino acid interaction in the trimer interface and lipids in the central cavity forming a pseudo-pore (Extended Data Fig. 3a-c). In the extracellular side of pseudo-pore, the N-terminal loop inserts into the pseudo-pore, stabilizing trimer formation (Extended Data Fig. 3d-e). In addition, the lipids of extracellular layer insert into the trimer interface formed by TM2 and TM3 and the TM4 of adjacent subunit (Extended Data Fig. 3f-g). Our purified protein contains the endogenous retinal (Fig. 2a-f), which is consistent with the electrophysiological experiments that *Hc*KCR1 can work without additional retinal supplementation. Therefore, our structure presents the natural functional state of *Hc*KCR1.

**Fig. 3.**
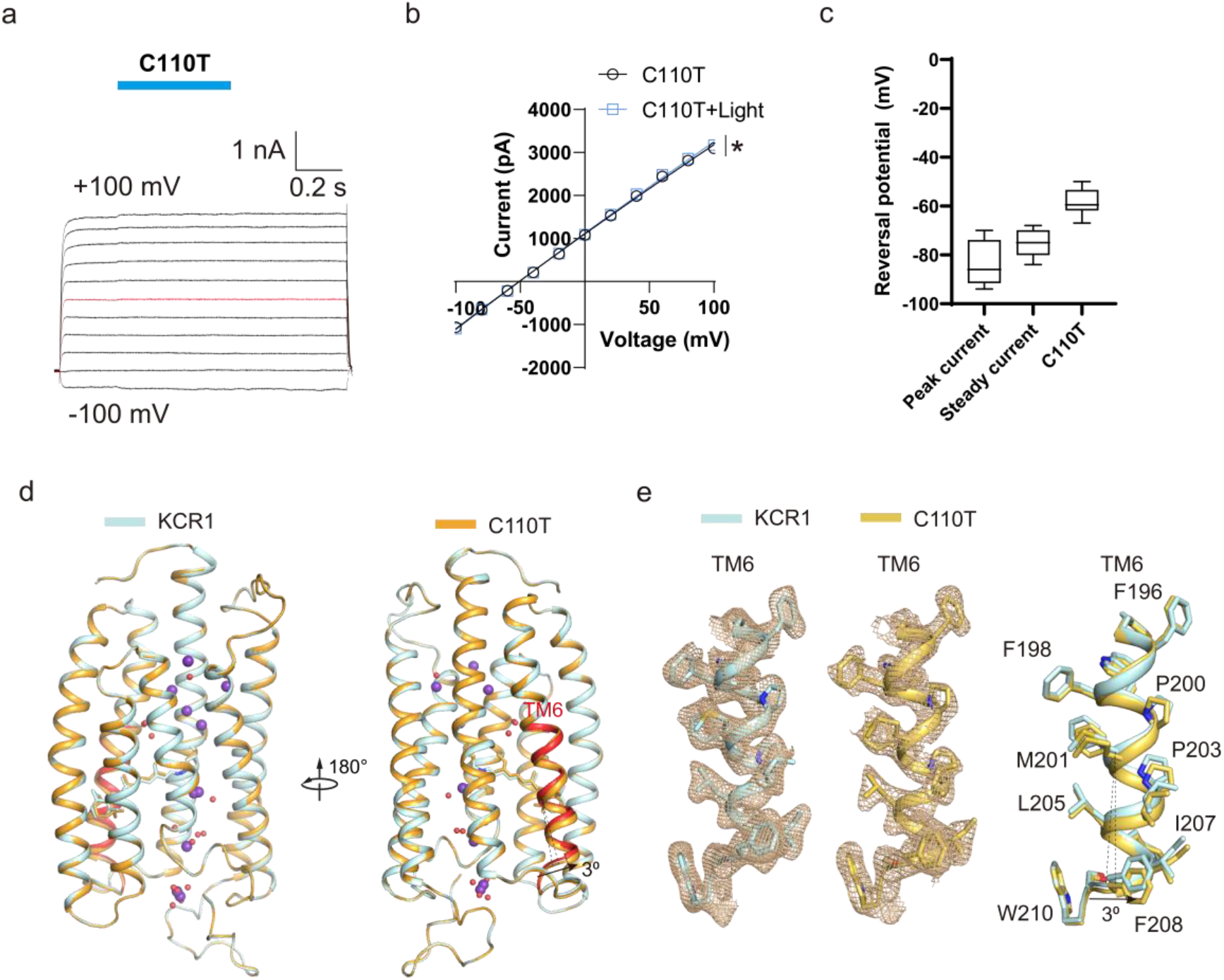
Conducting state of *Hc*KCR1-C110T. **a**, The representative physiological whole cell electrophysiological recordings from the HEK293T cell expressing *Hc*KCR1-C110T mutant with the voltage-pulse from-100 mV to +100 mV with a +20 mV step. Each step is stimulated by blue light and that is indicated by the blue line. The red trace stands for the current recorded at the holding potential of 0 mV. **b,** The current-voltage (I-V) plot of resting and light activated currents from (**a**). **c,** The reversal potential of peak currents and steady currents of wildtype *Hc*KCR1 and HcKCR1-C110T mutant. **d-e,** Structural comparison of wildtype *Hc*KCR1 and HcKCR1-C110T mutant. The red TM segment indicates that C110T mutant exhibits a slight extension toward the membrane side compared to the wildtype *Hc*KCR1.

### Structure of open state of *Hc*KCR1 (C110T)

Despite we can observe the unambiguous ions/water densities in each monomer, accurately discriminating potassium remains difficult. One of the most direct and effective alternatives to identify potassium is to obtain a high-resolution structure of the open state of *Hc*KCR1, because the continuous potassium must flow as the dominant ion in the ion permeation pathway. By using mutagenesis technology, we obtained an open state of *Hc*KCR1 (C110T). The C110T shows a spontaneous open state in the dark environment while light can enhance the ion flow (Fig. 3a-b). The reversal potential of the spontaneous current is-58.6 ± 3.7 mV (Fig. 3c) and the P_K_/P_Na_ permeability ratio is around 10, therefore it still retains the significant high potassium selectivity. Next, we purified C110T and solved the cryo-EM structure as the wild-type *Hc*KCR1 (Extended Data Fig. 4-5). The C110T is overall identical to the wild-type *Hc*KCR1 (Fig. 3d). One cryptic difference between wildtype *Hc*KCR1 and C110T is a slight extension of the transmembrane segment of extracellular side of TM6 (Fig. 3d-e). Carefully checking the retinal, we found the retinal in C110T appears to be hydrolyzed or broken, leaving a more than 3 Å gap that may allow dehydrated potassium to flow (Fig. 4a-f). Interestingly, we could observe that two tyrosine residues (Y103 and Y106) flip along the phenol ring together with the extracellular side of TM6 extension (Fig. 4g-i), which are mainly caused by the conformational changes of the retinal. Mutation of Y103 or Y106 to alanine results in fast inactivation of *Hc*KCR1 (Fig. 4j-l), which further suggest that Y103 and Y106 function critical roles in photoactivation. Therefore, the structure of C110T should present the open form of *Hc*KCR1. As the density of continuous ions/water occupying in the narrow lumen of C110T is essentially consistent with the wild-type cryo-EM map, both the ion densities in the wildtype map and C110T map present the continuous potassium flow (Extended Data Fig. 2 and 5).

**Fig. 4.**
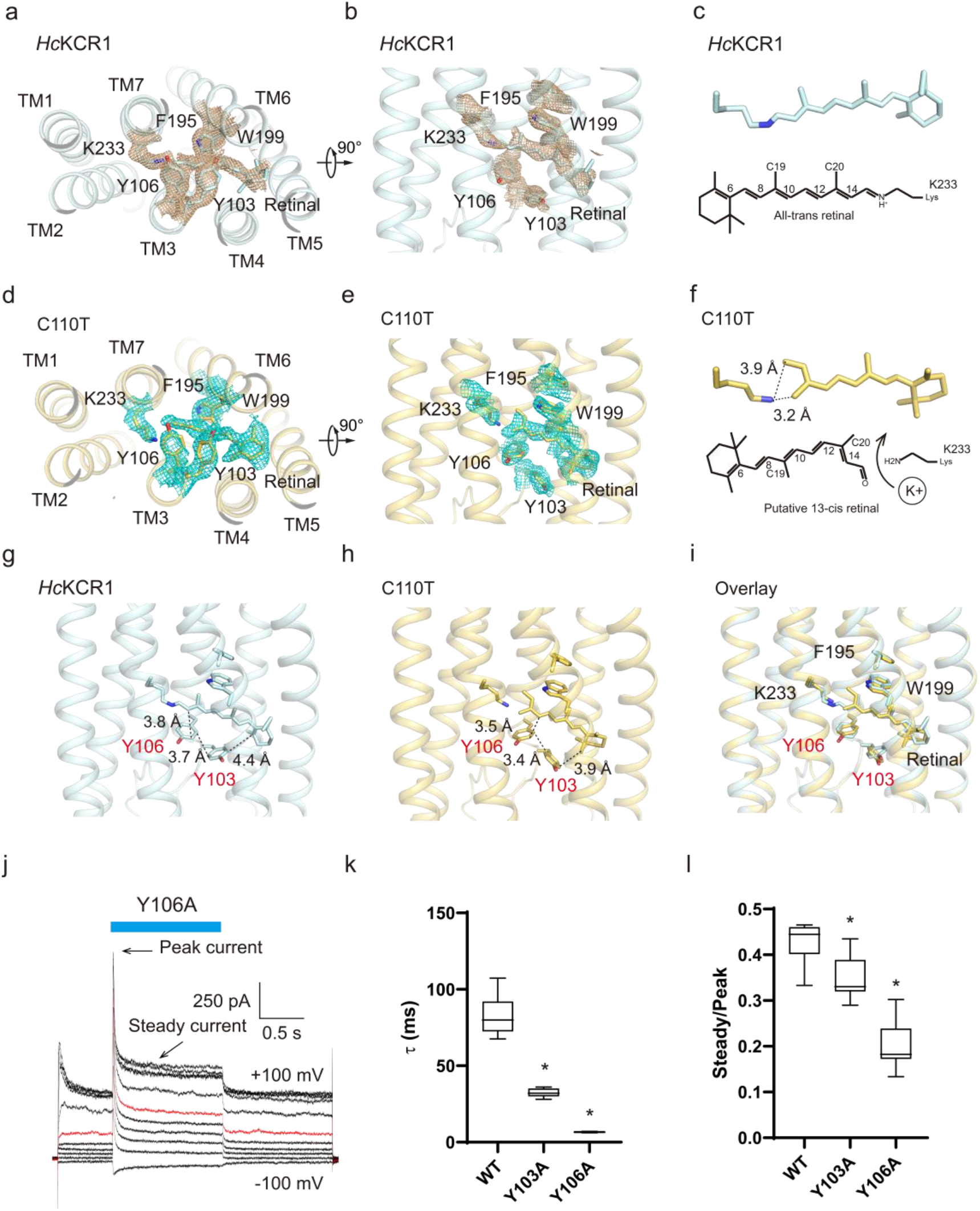
Conformational changes of the retinal and its coordination residues in the conducting form. **a-b**, The retinal binding pockets of *Hc*KCR1viewed from top **(a)** and side **(b)**. The coordination residues are marked in the figure. **c,** The potential Schiff-based linked all-trans retinal in the wildtype *Hc*KCR1. **d-e,** The retinal binding pockets of C110T mutant viewed from top **(d)** and side **(e)**. The coordination residues are marked in the figure. **f,** The potential hydrolyzed 13-cis retinal in the C110T mutant. **g-i,** The interaction between the Y103, Y106 and the retinal of wildtype *Hc*KCR1 **(g)**, C110T mutant **(h)** and the overlay view **(i)**. The side chain flip of Y103 and Y106 is coordinated with the conformational changes of the retinal. **j,** The representative physiological whole cell electrophysiological recordings from the HEK293T cell expressing *Hc*KCR1-Y106A mutant with the voltage-pulse from-100 mV to +100 mV with a +20 mV step. Each step is stimulated by blue light and that is indicated by the blue line. The red trace stands for the current recorded at the holding potential of 0 mV. The peak currents and inactivated steady currents are indicated. **k,** Inactivation decay time of wildtype, Y103A and Y106A mutants. **i,** The steady current and peak current ratio of wildtype, Y103A and Y106A mutants. The statistic data points are the Mean ± SEM (n is at least 5 cells for each variant). *, p < 0.05 by the student’s t-test.

**Fig. 5.**
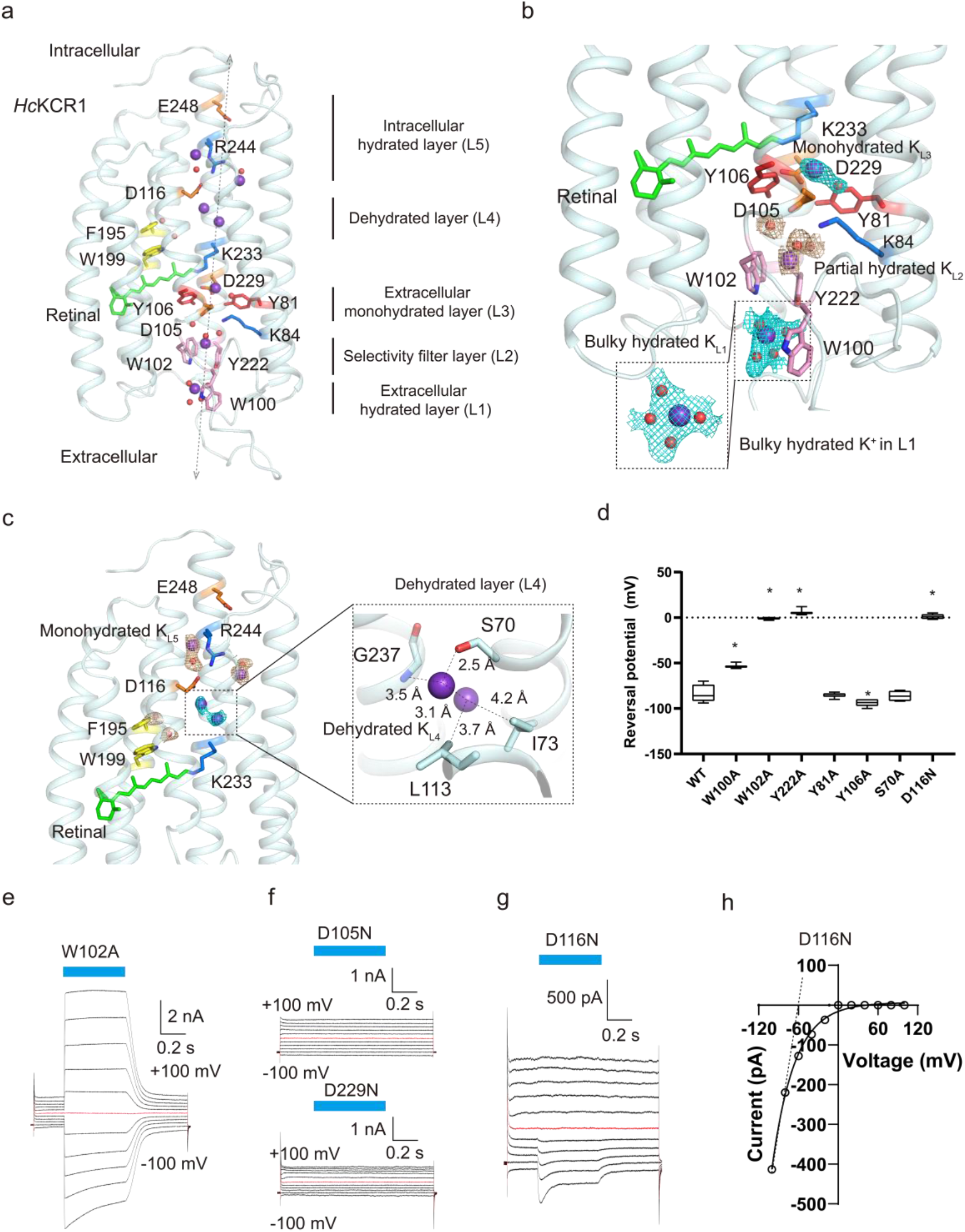
The continuous potassium permeation pathway of *Hc*KCR1. **a**, The water molecules are shown as smaller red/brown spheres and the potassium ion are shown as larger purple spheres. We divided the lumen from intracellular to extracellular side into five layers: the intracellular hydrated layer (L5, E248-R244-D116), which contains the hydrated potassium; the dehydrated layer (L4), containing two water molecules (F195-W199) and two continuous dehydrated potassium ion; the extracellular monohydrated layer (L3, K233-D229-D105-K84), which contains a monohydrate potassium ion; the selectivity filter layer (L2, W102-Y222), which contains at least one partial hydrated potassium ion through π-cation interactions. the extracellular hydrated layer (L1, W100), which contains the bulky hydrated potassium ion. **b,** The zoom view of extracellular lumen. We name the bulky hydrated potassium ion in the L1 as K_L1_, partial hydrated potassium ion in the L2 as K_L2_, monohydrated potassium ion in the L3 as K_L3_. **c,** The zoom view of intracellular lumen. We name the dehydrated potassium ions in the L4 as K_L4_, hydrated potassium ion in the L5 as K_L5_. The potential coordinated residues of the dehydrated potassium ions are marked in the figure. **d,** The reversal potential of peak currents of the indicated mutants. **e-g,** The representative physiological whole cell electrophysiological recordings from the HEK293T cell expressing W102A **(e)**, D105N, D229N **(f)** and D116N **(g)** mutants with the voltage-pulse from-100 mV to +100 mV with a +20 mV step. Each step is stimulated by blue light and that is indicated by the blue line. The red trace stands for the current recorded at the holding potential of 0 mV. **h,** The current-voltage (I-V) plot of light activated peak currents of D116N mutant from **(g)**. The dash lines indicate the virtual trendlines if the channels have no inward rectifier property. The farther away from this virtual dashed line, the greater the inward rectifier property.

### Identify key residues mediating potassium selectivity

Firstly, we divided the potential lumen region into five different layers (L1-L5) (Fig. 5a). The mutant (W100A) in L1 layer resulted in slightly reduced potassium selectivity, while the mutants (W102A and Y222A) in L2 layer led to largely reduced potassium selectivity. Additionally, the mutant (Y106A) of residues in L3 layer slightly increase the potassium selectivity. Surprisingly, the D116N mutant in L5 layer shows the typical strong voltage-dependent inward rectification manner that also eliminates the potassium selectivity (Fig. 5d-h). Therefore, the L1 and L2 layers, especially L2 layer, play vital roles in potassium selectivity. To further investigate the selectivity of amino acids for potassium, we mutate W102 and Y222 to 19 other amino acids, respectively, and found that the aromatic features of W102 and Y222 play a major role in potassium selectivity (Extended Data Fig. 6a-d). Similar partial functional results are reported during our submission^14, 15^.

**Fig. 6.**
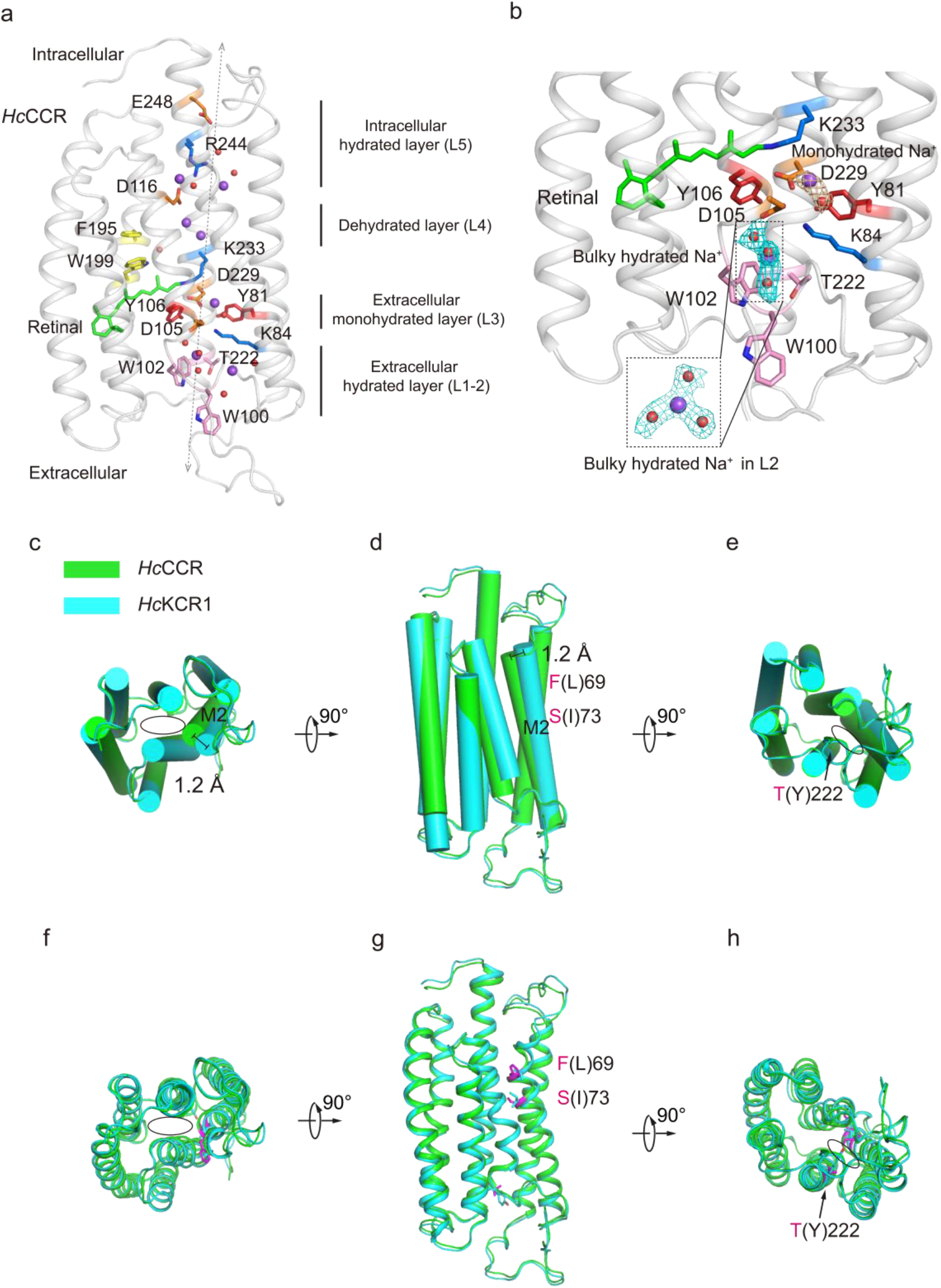
The lumen property and ion distributions of *Hc*CCR. **a-b**, The lumen property and ion distribution of *Hc*CCR are displayed like the *Hc*KCR1. The L2 of *Hc*CCR is occupied with the bulky sodium instead of the partial hydrated ion in the *Hc*KCR1. **c-e,** The cylinder display of top **(c)**, side **(d)** and bottom **(e)** views of *Hc*KCR1 (cyan) and *Hc*CCR (green). **f-h,** The canonical carton display of top **(f)**, side **(g)** and bottom **(h)** views of *Hc*KCR1 (cyan) and *Hc*CCR (green). The TM2 motion around 1.2 Å toward the lumen side. The motion starts from the S73 and F69. The cytosolic side blockage residue (Y222) is indicated in the **(e)** and **(h)**.

To address the unexpected role of D116N in potassium selectivity, we solved the cryo-EM structure of D116N (Extended Data Fig. 7a-c and 8) and found the only difference between wildtype *Hc*KCR1 and D116N mutant is the flip of R244, which is the potential interaction amino acid of D116. In the wildtype *Hc*KCR1, the side chain of R244 flips towards the side chain of D116 forming a salt-bridge-like structure (Extended Data Fig. 9a-f). In the D116N, the side chain of R244 flips towards E248 forming another salt-bridge-like structure. Therefore, the alternative formation of salt bridges between E248, R244 and D116 may convert hydrated potassium to dehydrated potassium. In addition, the E248Q mutant also exhibit the inwardly rectifying property, further support the dehydration-capable site (E248-R244-D116 pairs) mediating the voltage dependent dehydration process (Extended Data Fig. 9d). Therefore, the D116N may largely disrupt the dehydration capacity that prevents the ion efflux to the extracellular side. Consequently, it exhibits the non-selectivity like manner.

**Fig. 7.**
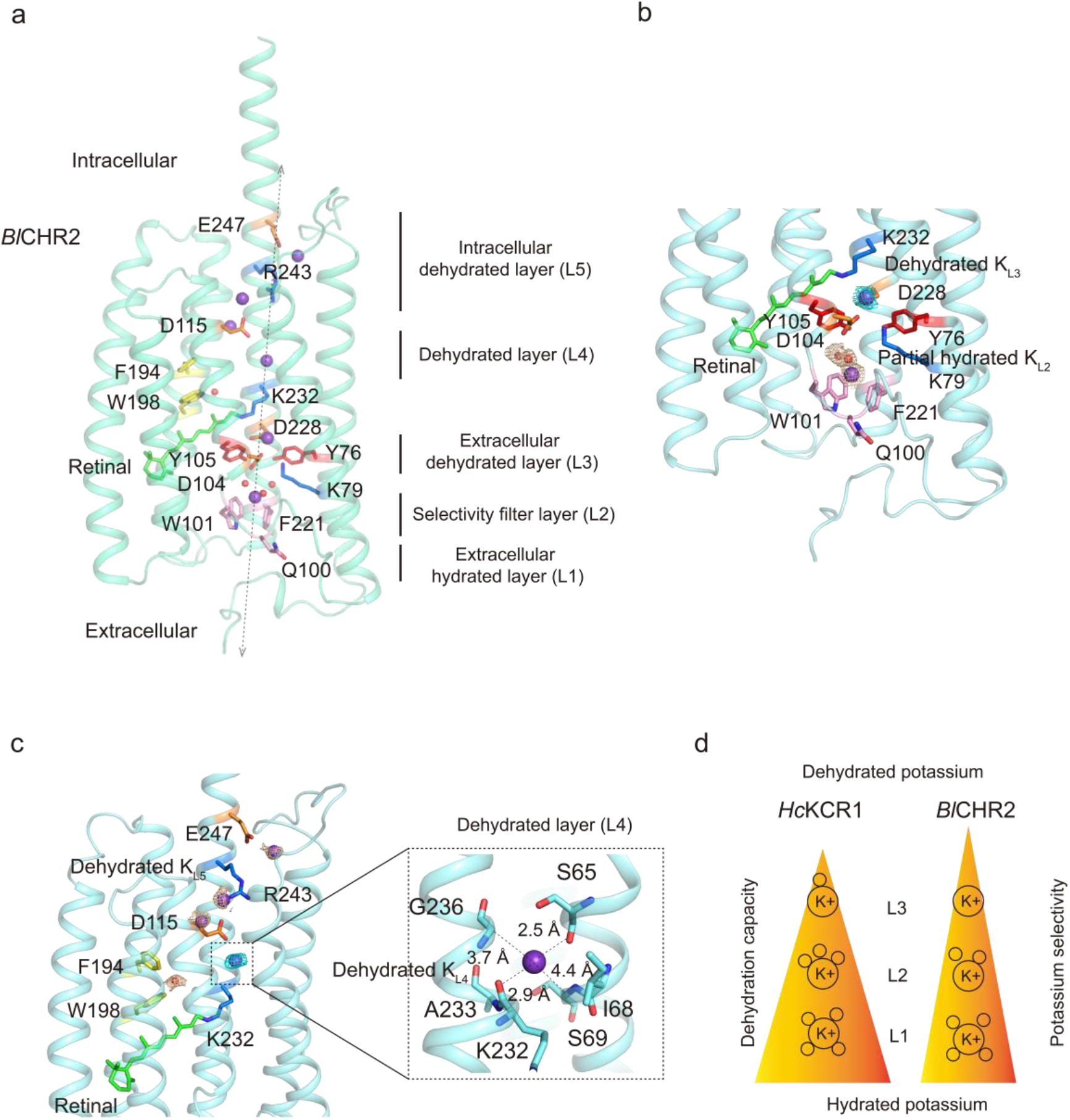
The lumen property and ion distributions of *Bl*CHR2. **a-c**, The lumen property and ion distribution of *Bl*CHR2 are displayed like the *Hc*KCR1. The L3 of *Bl*CHR2 is occupied with the dehydrated potassium instead of the monohydrated potassium in the *Hc*KCR1. **d,** The carton presents that the stepwise dehydration process of extracellular lumen and the stronger dehydration capacity, the higher potassium selectivity.

### Identify continuous potassium flow in the ion permeation pathway

The bulky potassium like density occupies in the L1, and we build it as the fully hydrated potassium (four water surround the potassium) and name it as K_L1_. Based on the strong ion density and the functional study, it is confident to build the potassium ion which directly interact with the aromatic side chain of W102 and Y222 in the L2 and we call it K_L2_. Meanwhile, at least 2-3 potential water molecules tightly interact with the K_L2_. However, depending on the geometry of the density, K_L2_ is not fully hydrated, but partially dehydrated. In the L3, based on the geometry of the ion density, it is confident to build as the monohydrated potassium (one potassium ion interacts with a water molecular) and we call it K_L3_ (Fig. 5b). In the L4, two continuous ion/water densities interact with the W199 and F195. The water molecule that interacts with W199 is conserved in other rhodopsins^22^. Therefore, we build these two continuous ion/water densities as two water molecules. Also, we build the two continuous dehydrated potassium ions, parallel to the two water molecules and we call them K_L4_. In the L5, we build two parallel monohydrated potassium separated by R244 and we call it K_L5_ (Fig. 5c). The hydrated potassium undergoes stepwise dehydration process along the extracellular lumen, which is likely mediated by the aromatic amino acids in the L2 and the alternative arrangement of two continuous pairs of acidic and alkali amino acids (K84-D105 and D229-K233) in the L3 (Fig. 5b). Mutations of D105 and D229 to asparagine eliminate the channel activity, further supporting the model (Fig. 5f). Additionally, the hydrated potassium also undergoes dehydration process along the intracellular lumen, which is mediated by the dehydration-capable site (E248-R244-D116 pairs) (Fig. 5c). Thus, the hydrated potassium undergoes dehydration process from both extracellular side and intracellular side of the lumen, and they all converge on the dehydrated layer L4.

### Narrower intracellular lumen chooses the smaller size of dehydrated sodium

Three residues reconstitution of *Hc*CCR to the corresponding residues of *Hc*KCRs convert the *Hc*CCR to potassium selelctivity^15^. Therefore, it offers a good material to study the ion selectivity of this special channelrhodopsin family. We solved the 2.37 Å resolution cryo-EM structure of *Hc*CCR and it allowed us to see clearly the ion/water density in the pore lumen (Extended Data Fig. 10a-c and 11). The overall structure of the *Hc*CCR is identical to the *Hc*KCR1, and they are both trimeric architectures (Extended Data Fig. 10c). As the Y222 is substituted by threonine in the L2, the ion/water in the L2 is the bulky hydrated sodium while the ion/water in the L3 is remain the monohydrated sodium (Fig. 6a-b). Therefore, the bulky hydrated ion is directly forced to conversion into the monohydrated ion by the conserved two continuous pairs of acidic and alkali amino acids (K84-D105 and D229-K233) in the L3 (Fig. 6b), which may not exclude the more difficult to dehydrate sodium ions.

Structural comparison of *Hc*CCR and *Hc*KCR1, we can observe around 1.2 Å contract of TM2 toward the pore, which make the intracellular lumen of *Hc*CCR narrower than the *Hc*KCR1 (Fig. 6c-e). Therefore, the narrower lumen may prevent the larger dehydrated potassium to flow but still allow the smaller dehydrated sodium to flow. In addition, the contract start from S73 and F69 (Fig. 4f-h), which is consistent with the substitution of S73 and F69 in the intracellular dehydrated lumen side as well as T222 in the L2 to the corresponding residues of *Hc*KCR1 convert the sodium selectivity of *Hc*CCR to potassium selectivity^15^. Taken together, narrower intracellular dehydrated ion occupied lumen and lose of the bulky hydrated sodium blockage in the L2 may allow for the sodium selectivity of *Hc*CCR.

### Higher dehydration capacity confers higher potassium ion selectivity

To further dissect the potassium selectivity mechanism, we solved two classes of cryo-EM structures of higher potassium selective channelrhodopsin (*B1*ChR2) from *Bilabrum sp*^14^ (Extended Data Fig. 12a-c and 13). The *B1*ChR2 also exhibit the trimeric architecture (Extended Data Fig. 12c-d). As the *B1*ChR2 have the additional long C-terminal region, one class of *B1*ChR2 presents an additional gate-like structure and another display the disordered C-terminal region (Extended Data Fig. 12d). Each class of the transmembrane region and the ion distribution of *B1*ChR2 is almost the same and they are identical to the *Hc*KCR1 (Extended Data Fig. 12d). Strikingly, the ion/water density in the L2 of *B1*ChR2 exhibits reduced water occupancy compared to *Hc*KCR1, while only one unambiguous density in the L3, thus, the K_L2_ presents less water surrounding configuration (further partially dehydrated potassium compared to K_L2_ of *Hc*KCR1) and K_L3_ presents the explicit dehydrated potassium configuration (Fig. 7a-b). Furthermore, in the L4 and L5, all ion/water densities exhibited single in *B1*ChR2 instead of two continuous or paired densities in *Hc*KCR1 (Fig. 7a and 7c), which suggested that the *B1*ChR2 adopt the continuous highly dehydrated potassium to confer the higher potassium ion selectivity (Fig. 7d). Therefore, the combination of the continuous ion distribution in the long and narrow lumen of *Hc*KCR1, *Hc*CCR and *B1*ChR2 largely suggests that the stepwise dehydration process in the extracellular side of *Hc*KCR1, *Hc*CCR and *B1*ChR2 play the dominant role in choosing the potassium ions that are more easily dehydrated and the intracellular side of *Hc*KCR1, *Hc*CCR and *B1*ChR2 play the vital role in choosing the right size of ions that are well occupied (Extended Data Fig. 14).

## Discussions

In *Hc*KCR1/*Hc*CCR/*B1*ChR2, the ion permeation pathway is a long and narrow hourglass shaped lumen across the whole cell membrane and gated by the covalently protonated Schiff base formed by the all-trans retinal chromophore and the conserved lysine^23^. These channelrhodopsins belong to the typical transporter like channels. The boundary between the ion channels and transporters is the speed of ion flow. Like the well-known CLCs (voltage dependent inwardly rectifying chloride channels or chloride transporters) superfamily, they share the sequence and structural similarity, but the channels and transporters have different functions^24^. The *Hc*KCR1/*Hc*CCR/*B1*ChR2 also can be divide into three distinct parts: extracellular hydrated, dehydrated, and intracellular hydrated parts, forming a sandwich-like structure that may safely guard the fast ion flow.

In the extracellular hydrated part of *Hc*KCR1/*B1*ChR2, there is a stepwise dehydrated potassium occupancy (L1-L2-L3) from extracellular side to the dehydrated layer. The aromatic amino acids in L2 layer likely act as a cache like filter role in producing a gradient of bulky hydrated potassium conversion to monohydrated (for *Hc*KCR1) and dehydrated (for *B1*ChR2) potassium, which may prevent the hydrated sodium ions that are not easily dehydrated. Thus, disrupting the L2 layer filter directly forces the conversion of bulky hydrated ions to monohydrated ions, thus eliminating potassium selectivity, as in the case of *Hc*CCR. Additionally, the higher dehydrated state of K_L3_ may confers the higher potassium selectivity. Interestingly, aromatic amino acids in KCRs interact with K_L2_ via cation-π interactions and the aromatic side chains are restrict. Some studies indicate that the affinity of cations with aromatic groups followed the order of K^+^ > Na^+^ > Li^+^^25^, which may give an additional explanation for the firm occupancy of K_L2_ in the L2. Also, in the intracellular hydrated part *Hc*KCR1, the conversion of hydrated potassium to dehydrated potassium is mediated by the potential dehydration-capable site (E248-R244-D116 pairs). It is worth noting that different sizes of dehydrating ions can be fine-tuned by different sizes of lumen, so the smaller size of intracellular lumen of *Hc*CCR allows the selection of smaller size of dehydrating sodium when the extracellular filter is disrupted.

The potassium selectivity mechanism of the traditional tetrameric potassium channels was extensively studied. It can be concluded as the restricted potassium selectivity filters, formed by the conserved sequence “T(S)VGY(F)G”, accommodate the continuous right size of dehydrated potassium for the fast flow^16^. Disruption one of the continuous potassium binding sites of potassium selectivity filters may result in loss of the potassium selectivity^26^. *Hc*KCR1 is strictly a potassium selective ion channel. The K_L2_ is highly bound to the aromatic amino acid in the L2 and the stepwise dehydrated potassium occupancy in the extracellular lumen reminiscence of the traditional potassium selectivity filter. One is stepwise dehydrated potassium occupancy in the long and narrow lumen and one is continuous dehydrated potassium occupancy in the long and narrow linear selectivity filter.

Due to the power of evolution, microbial rhodopsins can be ion channels, ion pumps, or enzymes^27^, while in the animal kingdom, the rhodopsins are mainly G-protein coupled receptors that act as master regulators of signal transduction cascades^28^. At the photoreaction level, light absorption induces photoisomerization from 11-cis to all-trans retinal (RET) resulting in movements in the transmembrane helix to form the active state in animal GPCR-type rhodopsins^29^. Usually, the retinal often forms a Schiff base with a lysin residue in rhodopsins, while in some vertebrate visual rhodopsins, the active state containing all-trans-retinal is thermally unstable, therefore, the Schiff base can be hydrolyzed and released from opsin after rhodopsin photo-bleaching^28^. The photoactivated state of *Hc*KCR1 is difficult to obtain, but we used mutagenesis to get a native open state in the dark environment. Surprisingly, based on the density of retinal, we found the retinal is likely to be hydrolyzed or broken, leaving a more than 3 Å gap that may allow dehydrated potassium flow. Although in some vertebrate visual rhodopsins, the Schiff base can be hydrolyzed and released from opsin after rhodopsin photo-bleaching, which provides a potential example to support the hydrolyzed or broken model, however, we recognized that the conformational changes in the retinal are limited and more studies need to be done in the future to figure out the entire photocycle coupled gating cycle.

## Methods

### Cloning and protein expression

The coding sequence for *Hc*KCR1, *Hc*CCR from *Hyphochytrium catenoides* and *B1*ChR2 from *Bilabrum sp* were cloned into a custom vector based on the BacMam expression backbone with an added C-terminal PreScission protease (PPX) cleavage site, linker sequence, enhanced GFP (EGFP), and a FLAG tag to generate the constructs KCRs/CCR-LE-LEVLFQGP-GGK-EGFP-GSG-DYKDDDDK for expression. Mutations of this study were introduced using inverse PCR. Constructs were transformed into DH10Bac *E.coli* to generate a bacmid according to the manufacturer’s instructions. Subsequent steps used *Spodopetera frugiperda* sf9 cells cultured in sf-900 II SFM medium (Thermo Fisher Scientific). Bacmid was transfected into adherent sf9 cells using the Cellfectin reagent (Gibco) to produce the P1 virus. P1 virus was used to infect sf9 cells in suspension at 1.5 million cells/mL at a multiplicity of infection (MOI) ∼0.1 to generate the P2 virus. Infection was monitored by fluorescence and the P2 virus was harvested 48-72 hours post-infection. P3 virus was generated in a similar manner. P3 viral stock was then used to infect Expi-HEK293F cells at 2.5 million cells/mL at an MOI ∼2-5 for large-scale protein expression. Cells were harvested by centrifugation at 4000×g for 15 min, flash-frozen in liquid nitrogen, and stored at -80°C.

### Protein purification and EM sample preparation

Cells from 1 L of culture were thawed and resuspended in 100 mL of lysis buffer (20 mM Tris-HCl, 150 mM KCl pH 8.0). Cells were lysed by sonication and membranes were pelleted by centrifugation at 40,000 r.p.m. for 40 min in a Ti45 rotor. The supernatant was loaded onto anti-FLAG resin (Genscript) by gravity flow. The resin was further washed with 10 column volumes of wash buffer (lysis buffer plus 0.02% LMNG), and protein was eluted with an elution buffer (wash buffer plus 230 μg/ml FLAG peptide). The C-terminal GFP tag of eluted protein was removed by HRV3C protease cleavage for 2 h at 4°C. The protein was further concentrated by a 100-kDa cutoff concentrator (Millipore) and loaded onto a Superose 6 increase 10/300 column (GE Healthcare) running in lysis buffer plus 0.03% digitonin. Peak fractions were combined and concentrated to around 10 mg/ml for cryo-EM sample preparation. The protein samples were cleared by centrifugation at 35,000 r.p.m. for 30 min at 4°C prior to grid preparation. A 3-μl drop of protein was applied to fresh glow discharged Holey Carbon, 300 mesh R1.2/1.3 gold grids (Quantifoil). Samples were plunge frozen in liquid nitrogen-cooled liquid ethane using an FEI Vitrobot Mark IV (Thermo Fisher Scientific) at 8°C, 100% humidity, 3 blot force, ∼5 s wait time, and 6 s blot time. All the samples were prepared under white light conditions.

### Cryo-EM data acquisition, processing, and model building

Cryo-EM data were collected on a Titan Krios microscope (FEI) equipped with a cesium corrector operated at 300 kV. Movie stacks were automatically acquired with EPU software on a Thermo Fisher Falcon4i detector with pixel size 0.35 or 0.45 or 0.57 Å at the object plane and with defocus ranging from-0.9 μm to-1.2 μm and GIF Quantum energy filter with a 5 eV slit width. The total exposure dose was 40 e^-^ Å^-2^. Acquisition parameters were summarized in Table S1. Data processings were carried out with the cryoSPARC v4 suite and were summarized in Extended figures. The atomic models of monomers were built in Coot based on an initial model (PDB: 8H22). The models were then manually adjusted in Coot. Trimeric models were obtained subsequently by applying a symmetry operation on the monomer. These trimeric models were refined using Phenix.real_space_refine with secondary structure restraints and Coot iteratively. For validation, FSC curves were calculated between the final models and EM maps. Figures were prepared using PyMOL and Chimera.

### Whole-cell patch clamp recording from HEK293 cells

The HEK293T cells from the American Type Culture Collection (ATCC) were cultured on coverslips placed in a 12-well plate containing DMEM/F12 medium (Gibco) supplemented with 10% fetal bovine serum (FBS). The cells in each well were transiently transfected with 1 μg DNA plasmid using 3μg linear polyethyleneimine (PEI, MW 25000, Polysciences) according to the manufacturer’s instructions. After 12-20 h, the coverslips were transferred to a recording chamber containing the external solution (10 mM HEPES-Na pH 7.4, 150 mM NaCl, 5 mM glucose, 2 mM MgCl_2,_ and 1 mM CaCl_2_). Borosilicate micropipettes (OD 1.5 mm, ID 0.86 mm, Sutter) were pulled and fire polished to 4-6 MΩ resistance. The pipette solution was 10 mM HEPES-Na pH 7.4, 150 mM KCl, and 5 mM EGTA. The bath solution was 10 mM HEPES-Na pH 7.4, 150 mM NaCl, 5 mM glucose, 2 mM MgCl_2,_ and 1 mM CaCl_2_. All measurements were carried out at room temperature (∼25°C) in the dark environment using an Axopatch 700B amplifier, a Digidata 1550 digitizer, and pCLAMP software (Molecular Devices). A blue light source is generated by the laser and controlled by pCLAMP software. The patches were held at-60 mV and the recordings were low-pass filtered at 1 kHz and sampled at 20 kHz. All current-voltage curves (IV dependencies) were corrected for liquid junction potentials (LJP) calculated using the ClampEx built-in LJP calculator. Statistical analyses were performed using ClampFit and GraphPad Prism.

**Extended Data Fig. 1.**
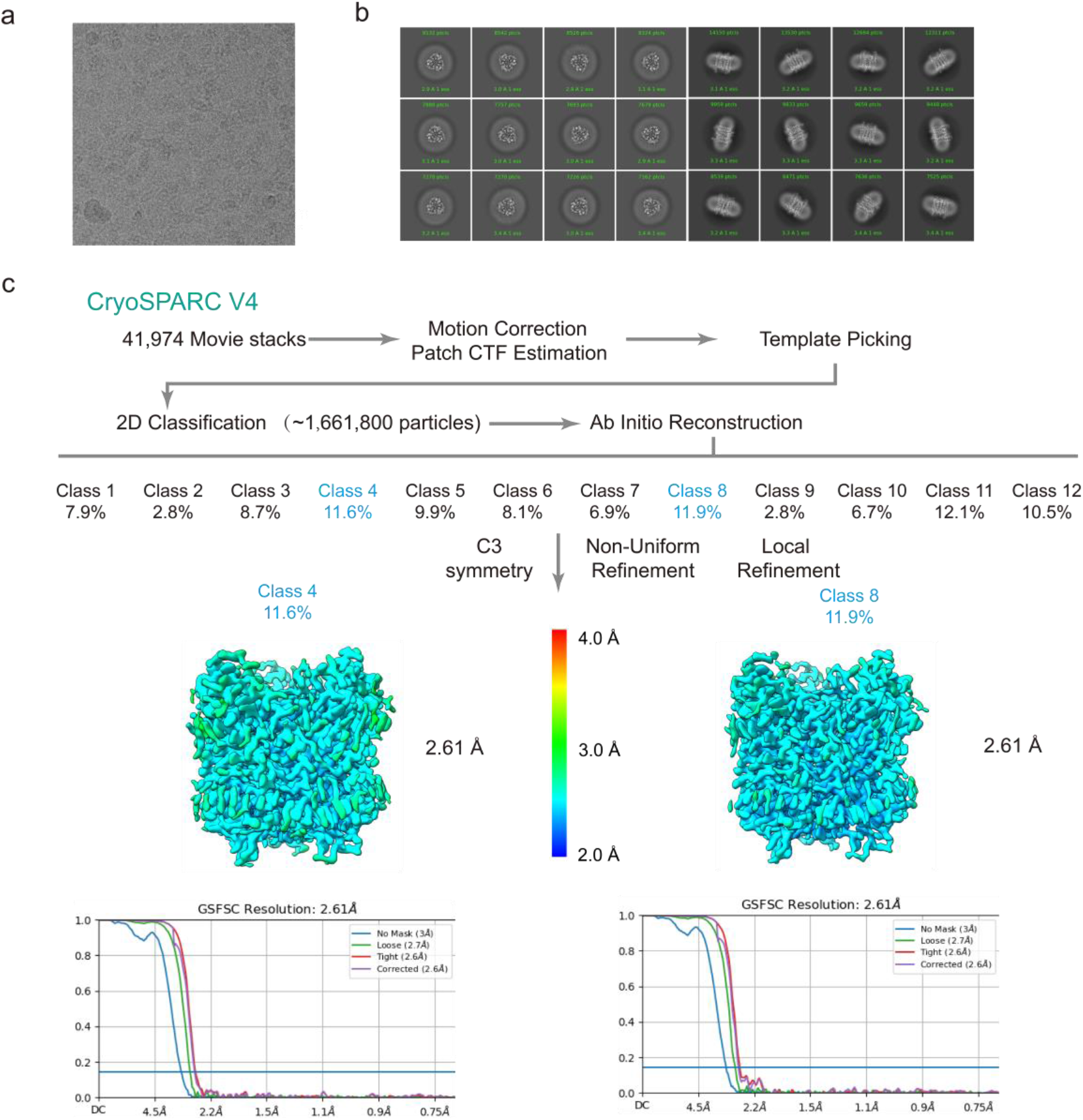
Single-particle Cryo-EM reconstructions of the *Hc*KCR1. **a**, A representative raw micrograph of the *Hc*KCR1. **b**, The selected 2D class averages. **c**, Summary of image processing for *Hc*KCR1 dataset with C3 symmetry.

**Extended Data Fig. 2.**
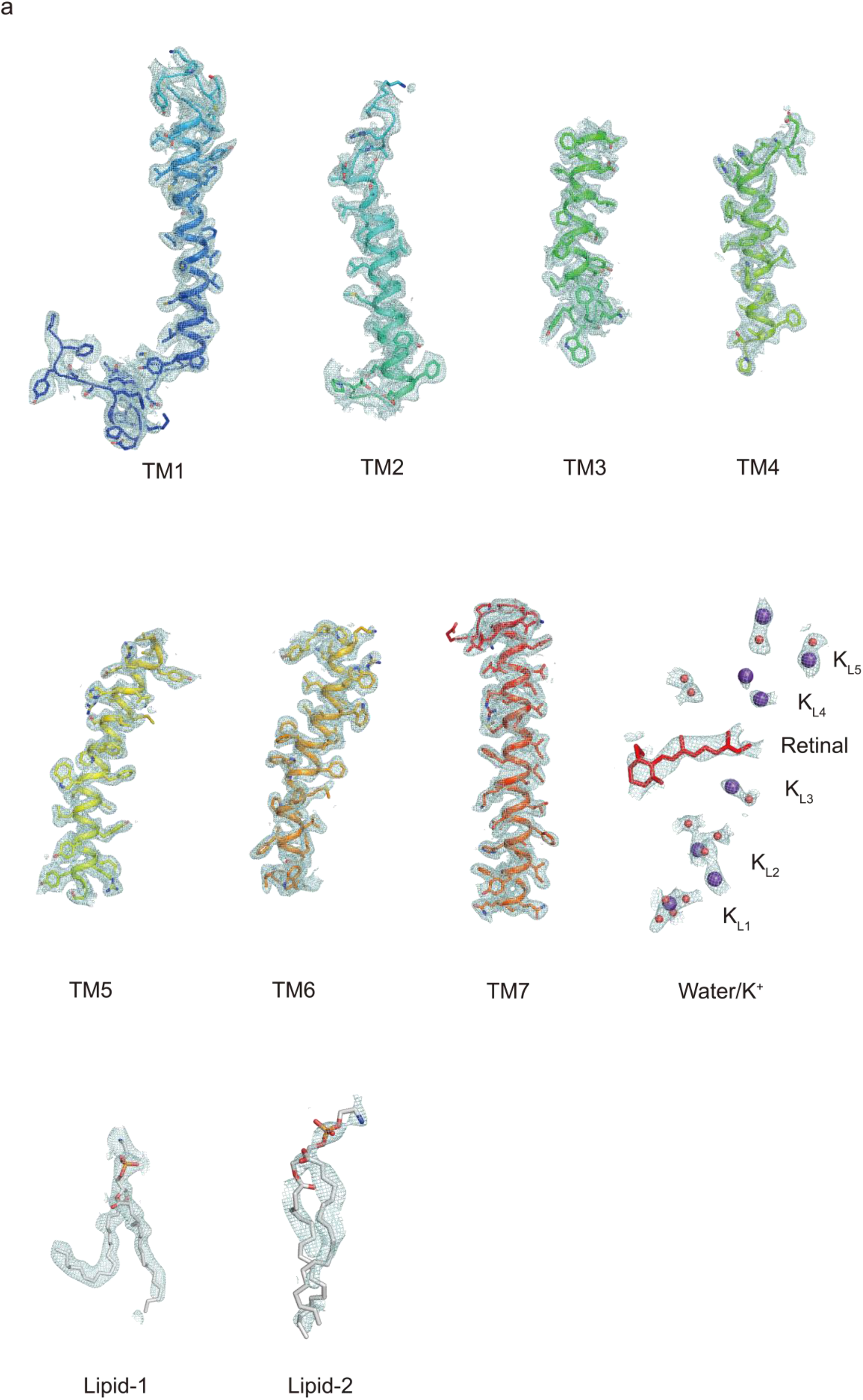
Cryo-EM density of *Hc*KCR1 in potassium environment. **a**, The cryo-EM density of the residues, ion/waters, lipids and the endogenous retinal of wildtype *Hc*KCR1 in potassium environment.

**Extended Data Fig. 3.**
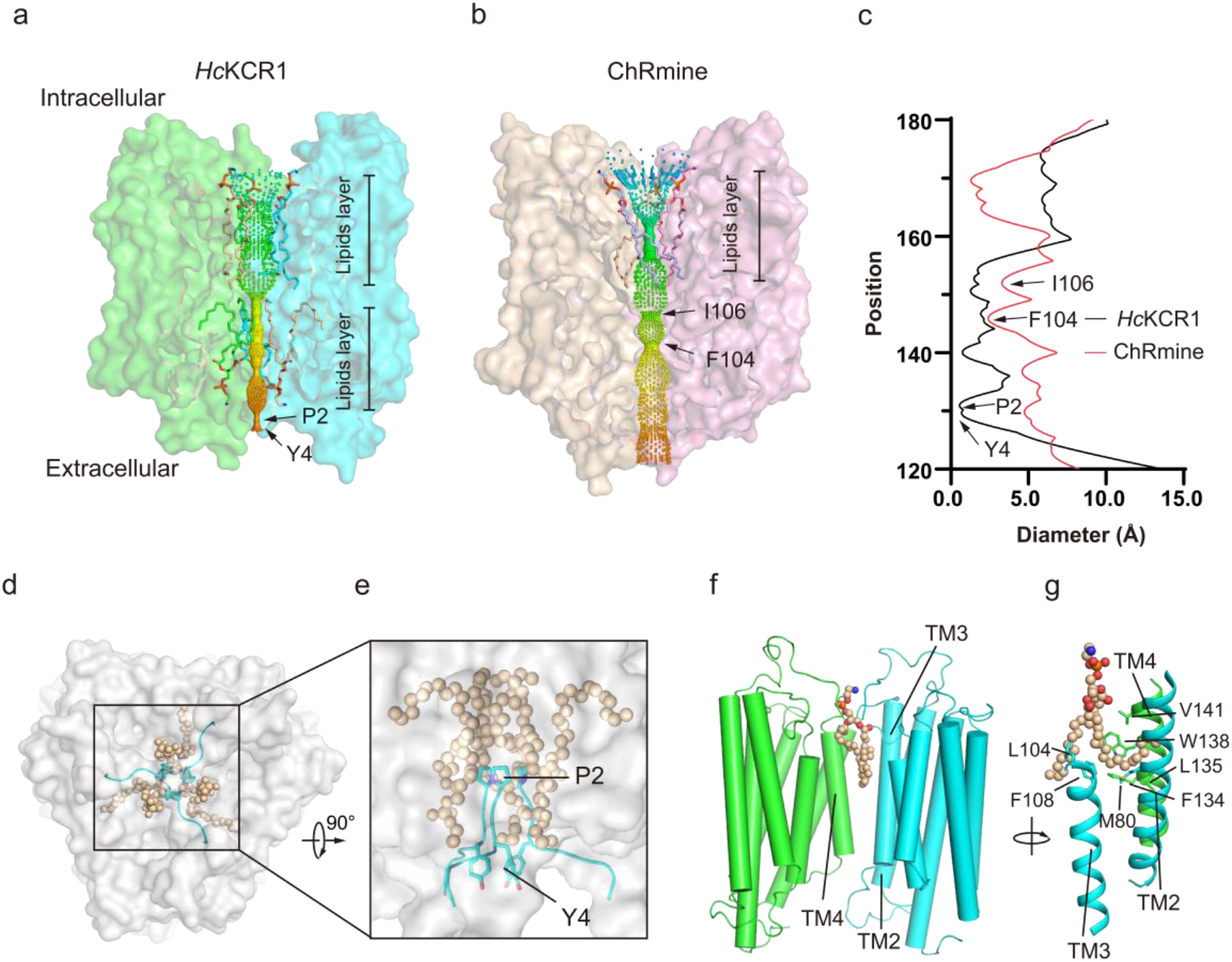
Trimeric organization of *Hc*KCR1. **a-b**, The *Hc*KCR1 pseudo-pore with two lipid layers **(a)** and the ChRmine pseudo-pore with one lipid layer **(b)** against the surface of two channel subunits (the front subunit is not displayed) viewed from the membrane plane. The restriction site, P2 and Y4 in *Hc*KCR1, I106 and F104 in ChRmine, are indicated by arrows. **c,** Pseudo-pore radius of *Hc*KCR1 and ChRmine as a function of distance from the intracellular side is shown. **d-c,** Constrictions of the pseudo-pore formed by lipids and residues (Y4 and P2) viewed form the extracellular side (**d**) and the membrane plane (**e**). **f-g,** The pseudo-pore lipids binding site in *Hc*KCR1 viewed from two angle. The key amino acids residues are indicated by arrows.

**Extended Data Fig. 4.**
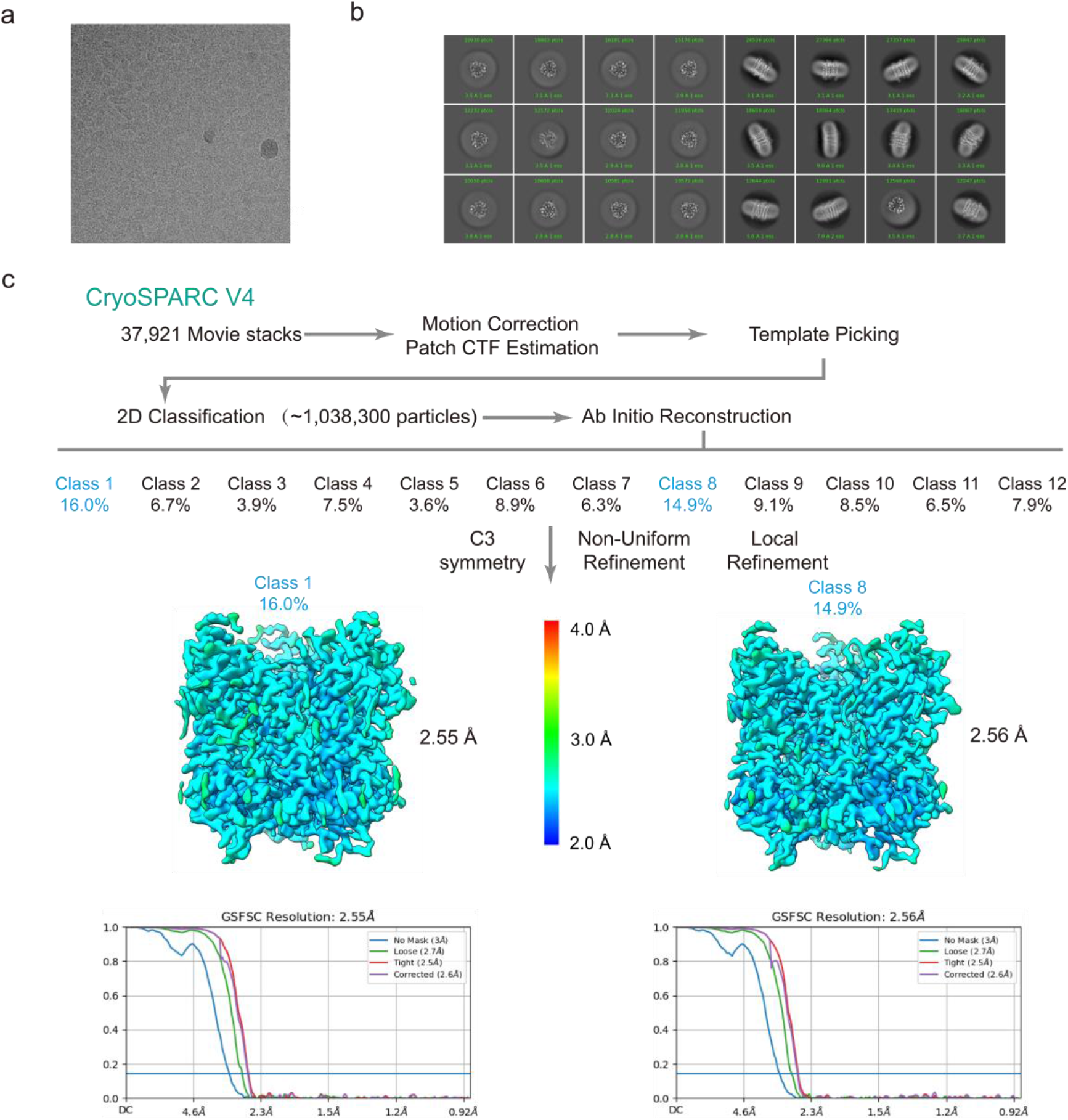
Single-particle Cryo-EM reconstructions of the *Hc*KCR1-C110T mutant. **a**, A representative raw micrograph of the *Hc*KCR1-C110T mutant. **b**, The selected 2D class averages. **c**, Summary of image processing for *Hc*KCR1-C110T mutant dataset with C3 symmetry.

**Extended Data Fig. 5.**
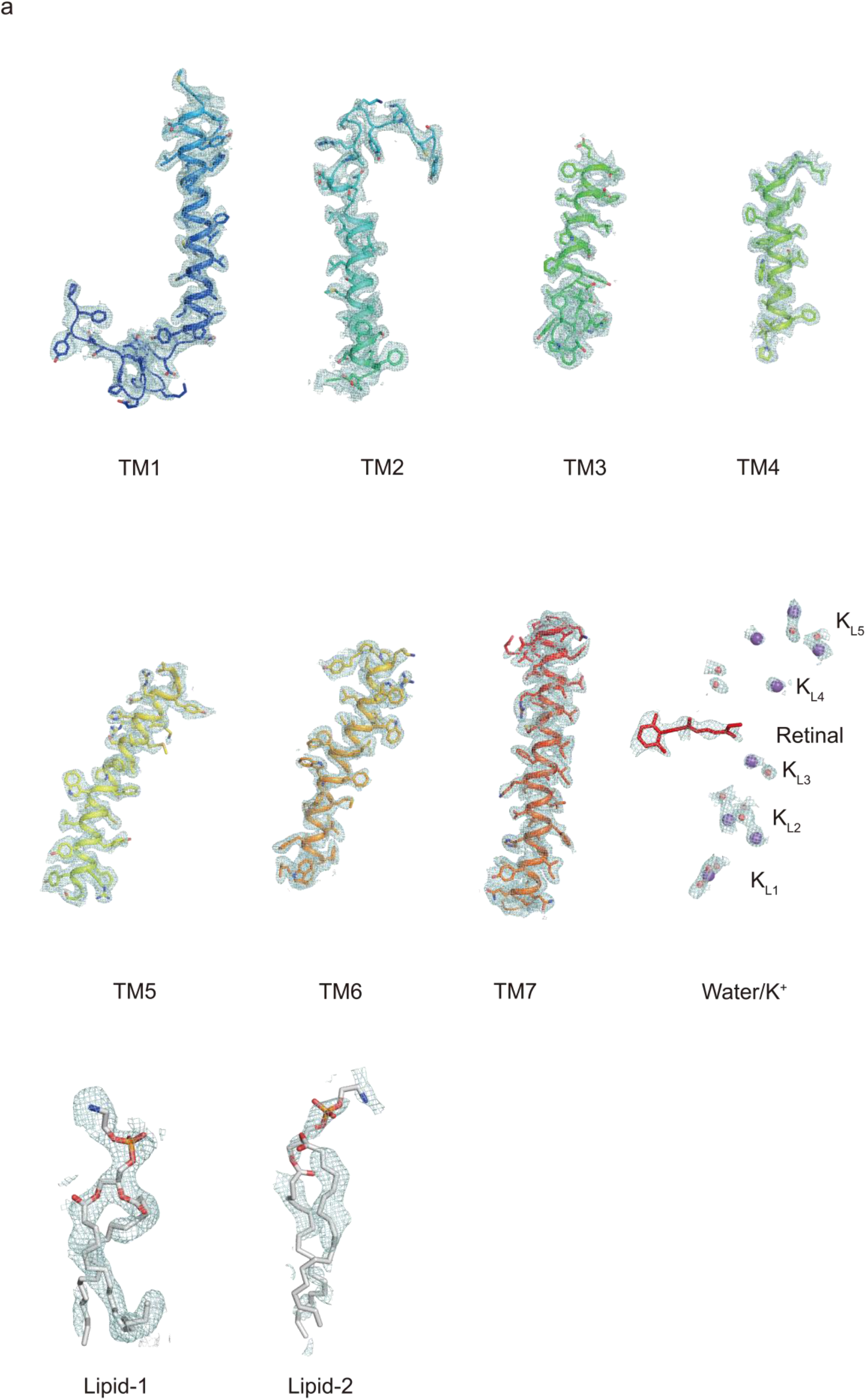
Cryo-EM density of *Hc*KCR1-C110T mutant in potassium environment. **a**, The cryo-EM density of the residues, ion/waters, lipids and the endogenous retinal of *Hc*KCR1-C110T mutant in potassium environment.

**Extended Data Fig. 6.**
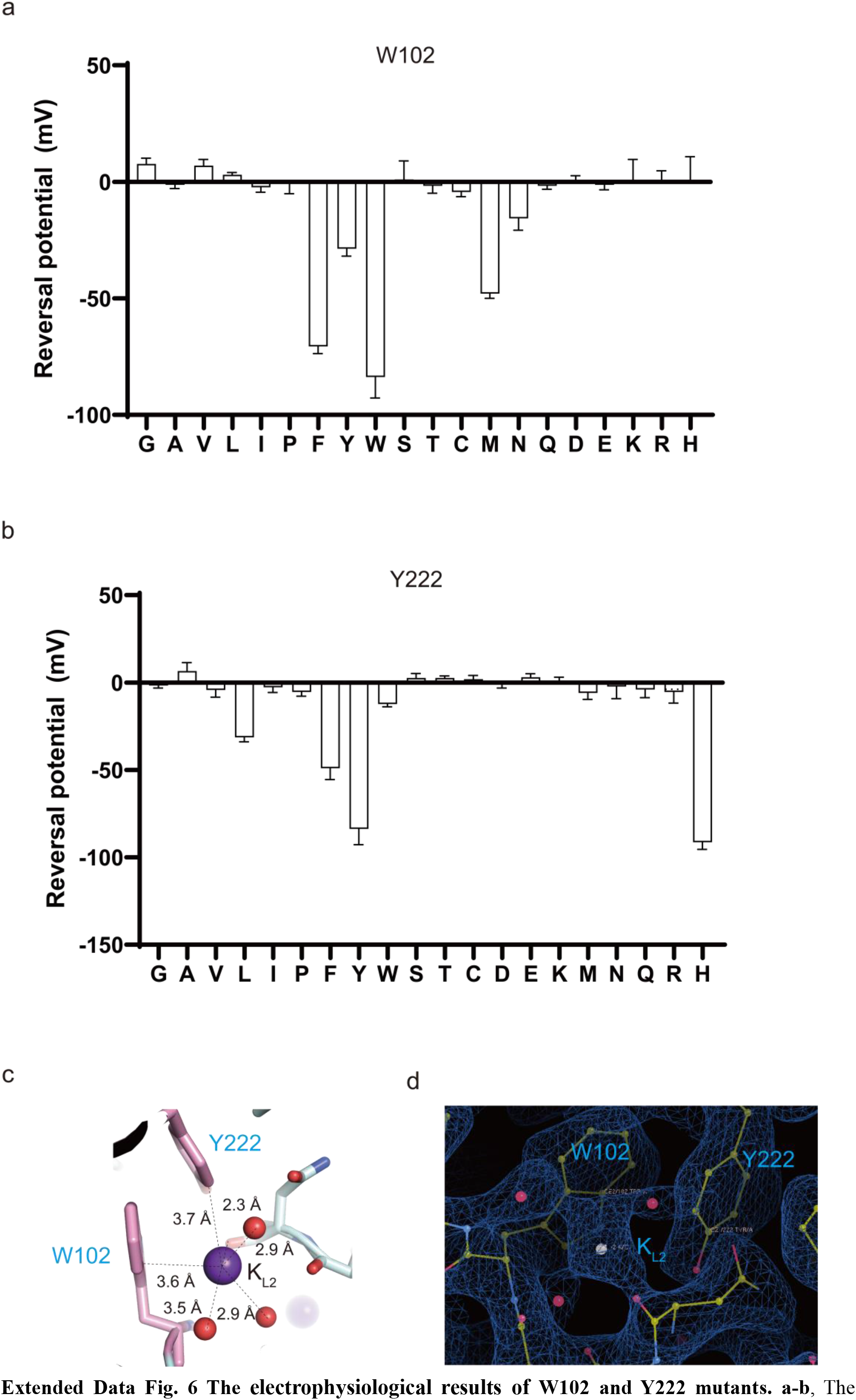
The electrophysiological results of W102 and Y222 mutants. **a-b**, The reversal potentials of the 19 amino acids substitutions of W102 and Y222. **c-d**, The W102 and Y222 interact with K_L2_ through the potential π-cation interactions.

**Extended Data Fig. 7.**
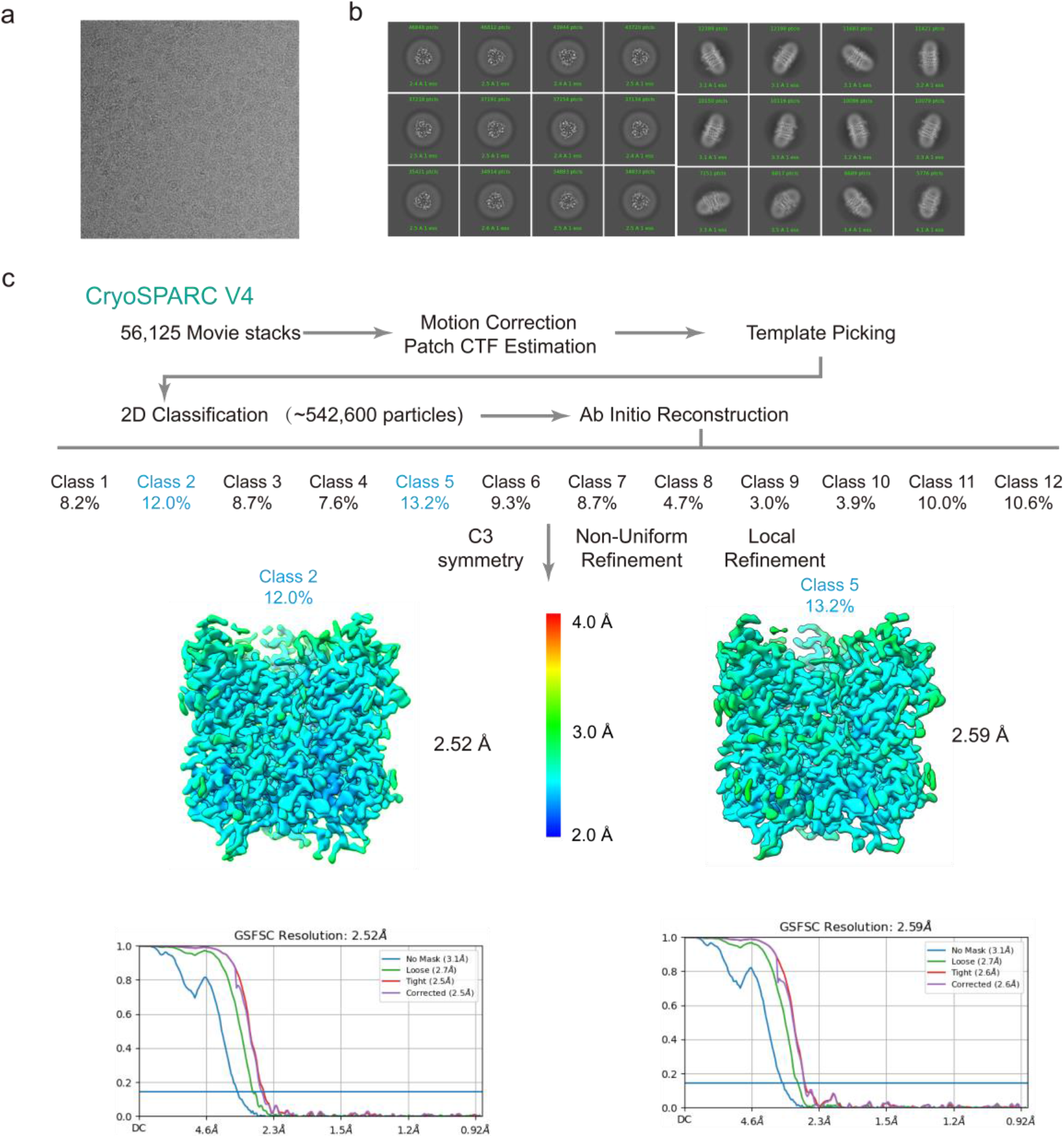
Single-particle Cryo-EM reconstructions of the *Hc*KCR1-D116N mutant. **a**, A representative raw micrograph of the *Hc*KCR1-D116N mutant. **b**, The selected 2D class averages. **c**, Summary of image processing for *Hc*KCR1-D116N mutant dataset with C3 symmetry.

**Extended Data Fig. 8.**
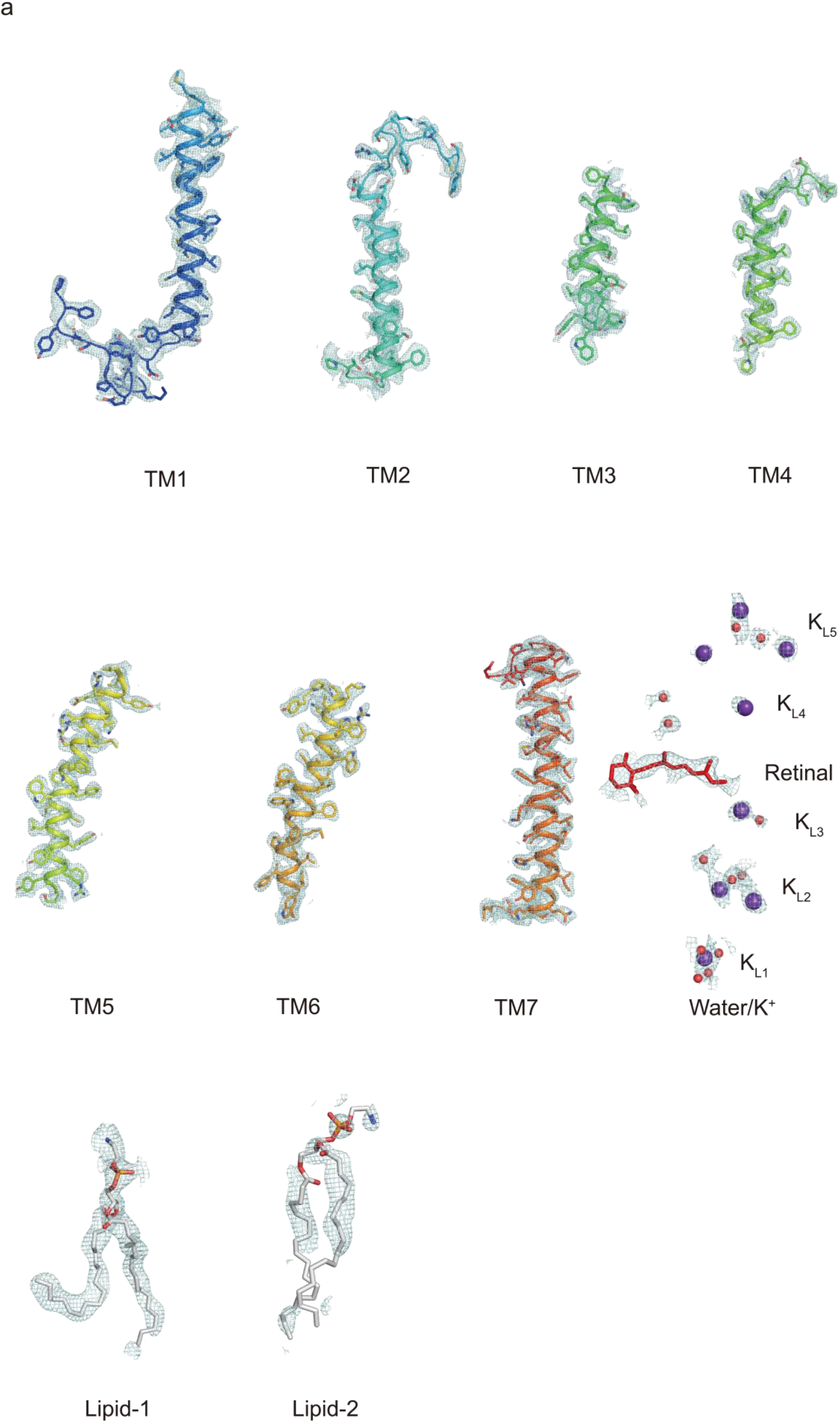
Cryo-EM density of *Hc*KCR1-D116N mutant in potassium environment. **a**, The cryo-EM density of the residues, ion/waters, lipids and the endogenous retinal of *Hc*KCR1-D116N mutant in potassium environment.

**Extended Data Fig. 9.**
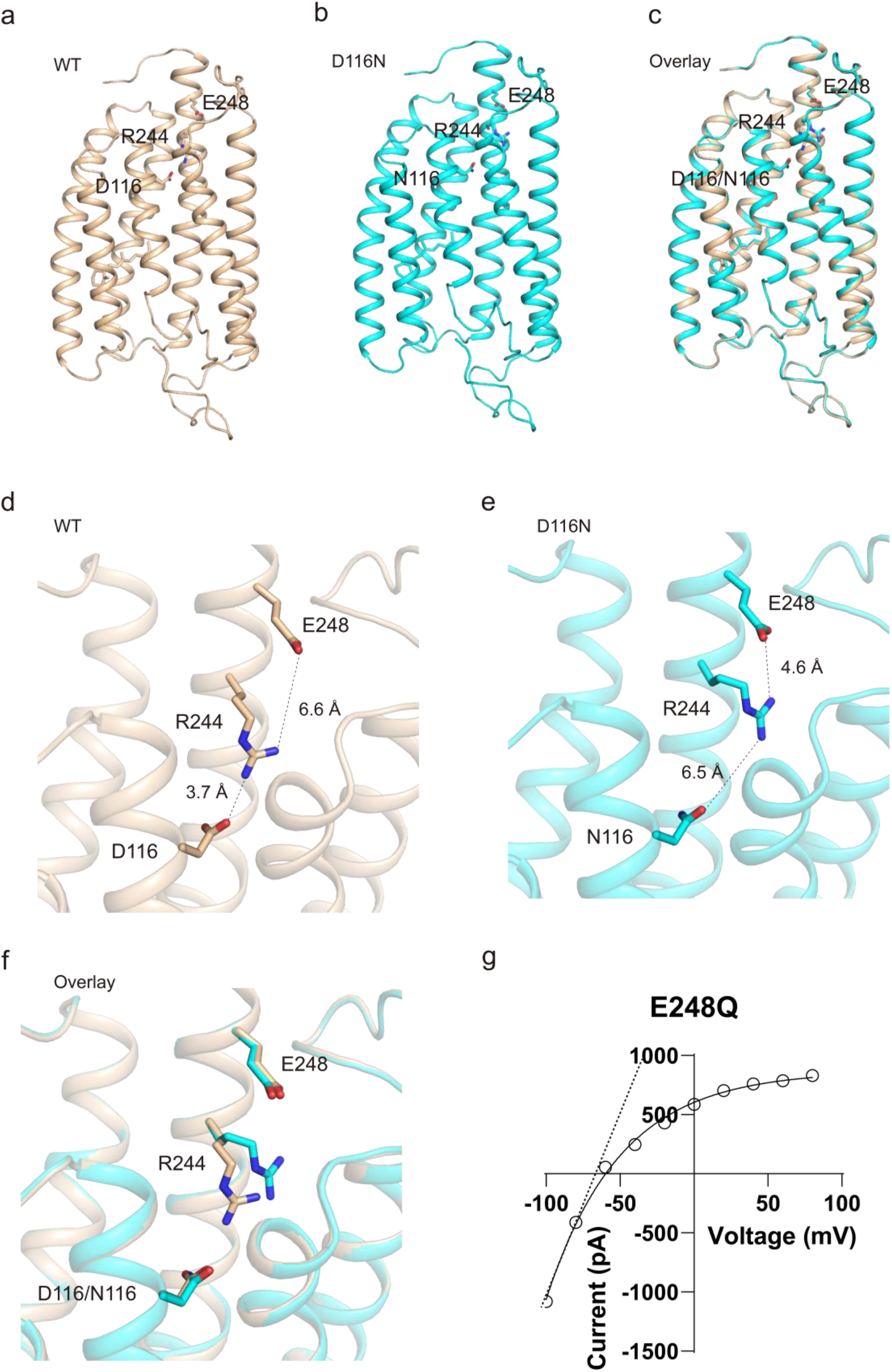
Structural comparison of wildtype *Hc*KCR1 and D116N mutant. **a-c**, The overall structural comparison of wildtype *Hc*KCR1 and D116N mutant. **d-f**, The intracellular dehydration-capable site (E248-R244-D116) of wildtype *Hc*KCR1 **(d)** and D116N **(e)** mutant. **(f)** is the overlay present of **(d)** and **(e). g**, The representative I-V plot of light activated peak currents of E248Q mutants at physiological whole cell configurations. The dash lines indicate the virtual trendlines if the channels have no inward rectifier property. The farther away from this virtual dashed line, the greater the inward rectifier property. Therefore, E248Q shows the strong inward rectifier property.

**Extended Data Fig. 10.**
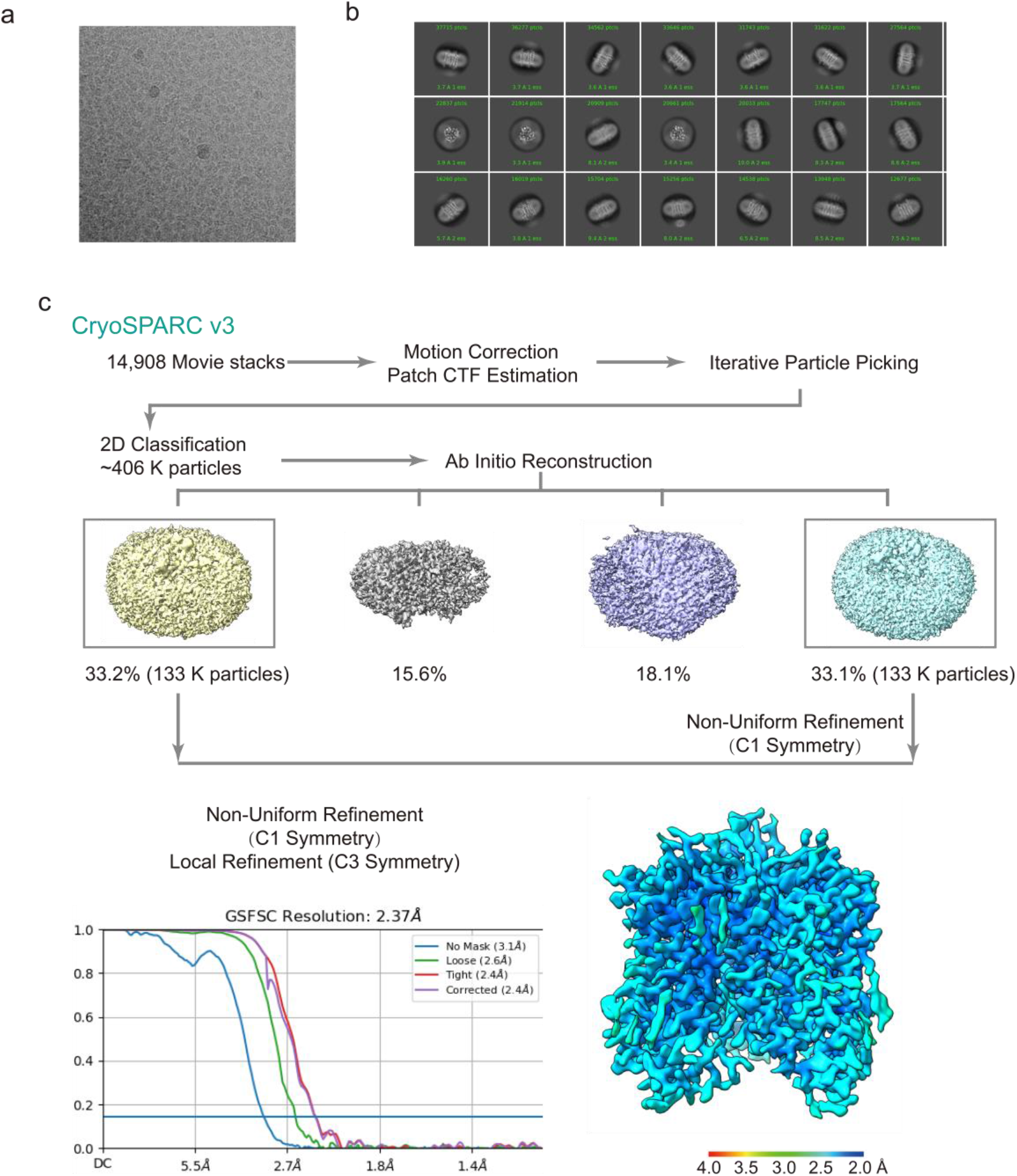
Single-particle Cryo-EM reconstructions of the *Hc*CCR. **a**, A representative raw micrograph of the *Hc*CCR. **b**, The selected 2D class averages. **c**, Summary of image processing for *Hc*CCR dataset with C3 symmetry.

**Extended Data Fig. 11.**
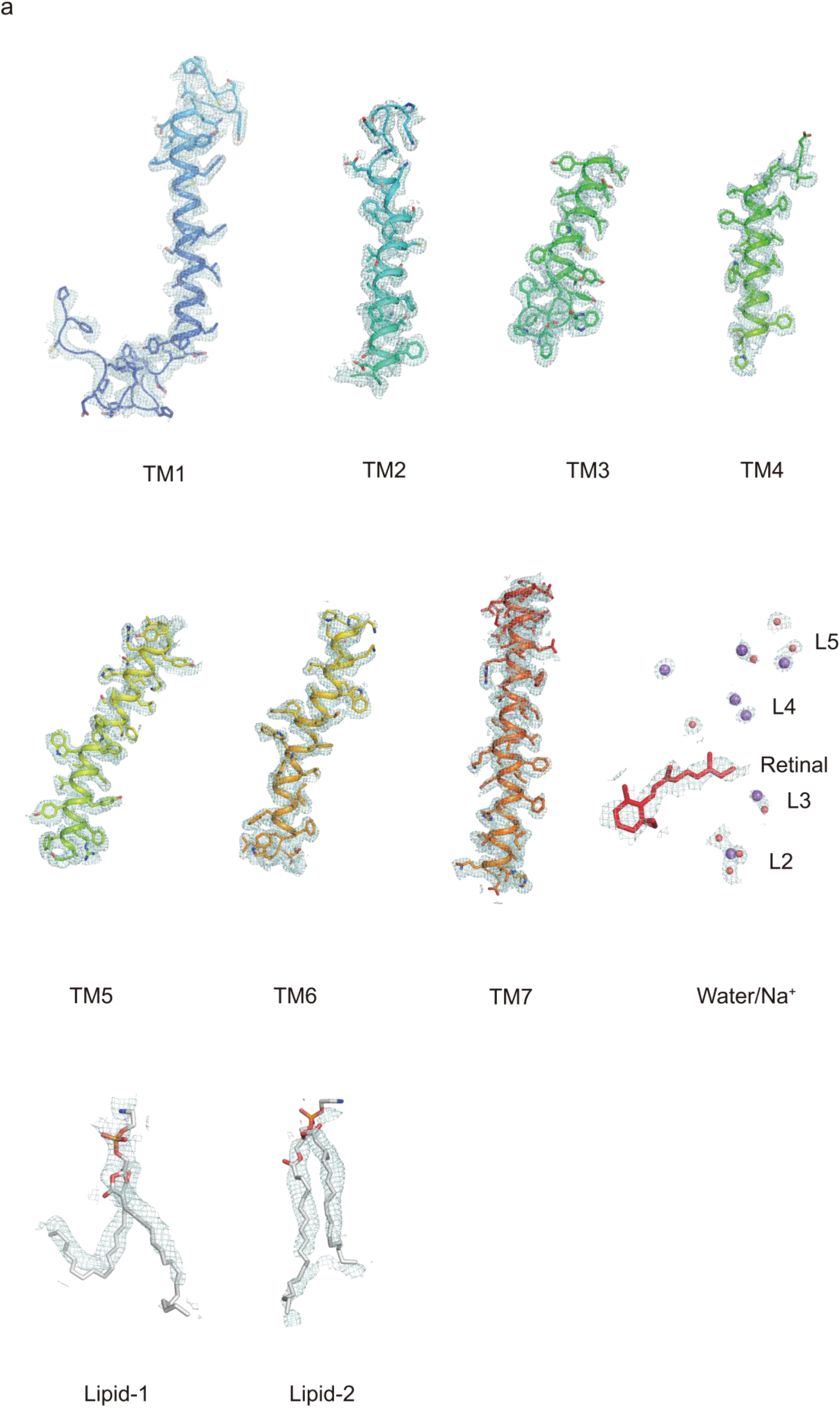
Cryo-EM density of *Hc*CCR in sodium environment. **a**, The cryo-EM density of the residues, ion/waters, lipids and the endogenous retinal of *Hc*CCR in sodium environment.

**Extended Data Fig. 12.**
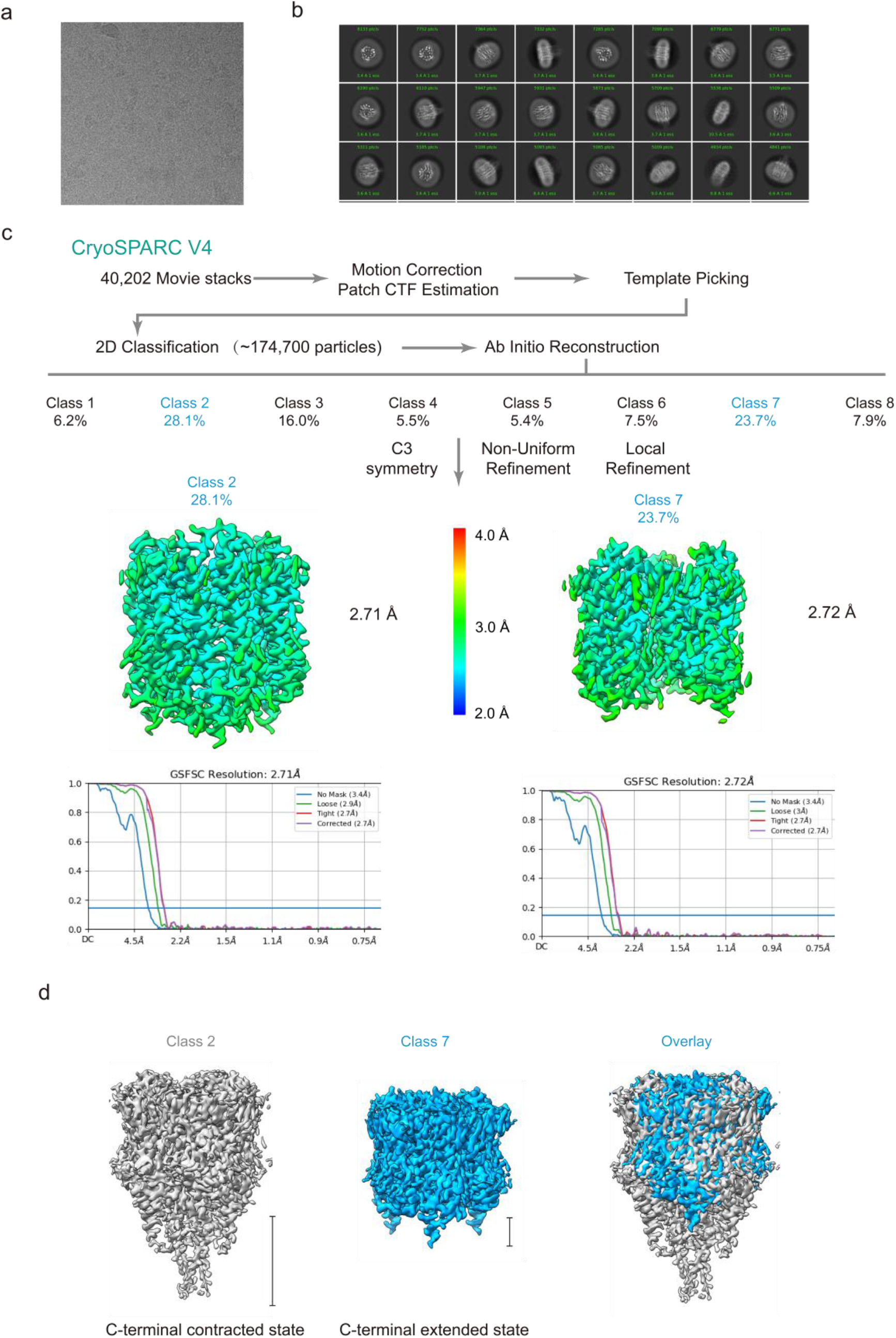
Single-particle Cryo-EM reconstructions of the *Bl*CHR2. **a**, A representative raw micrograph of the *Bl*CHR2. **b**, The selected 2D class averages. **c**, Summary of image processing for *Bl*CHR2 dataset with C3 symmetry.

**Extended Data Fig. 13.**
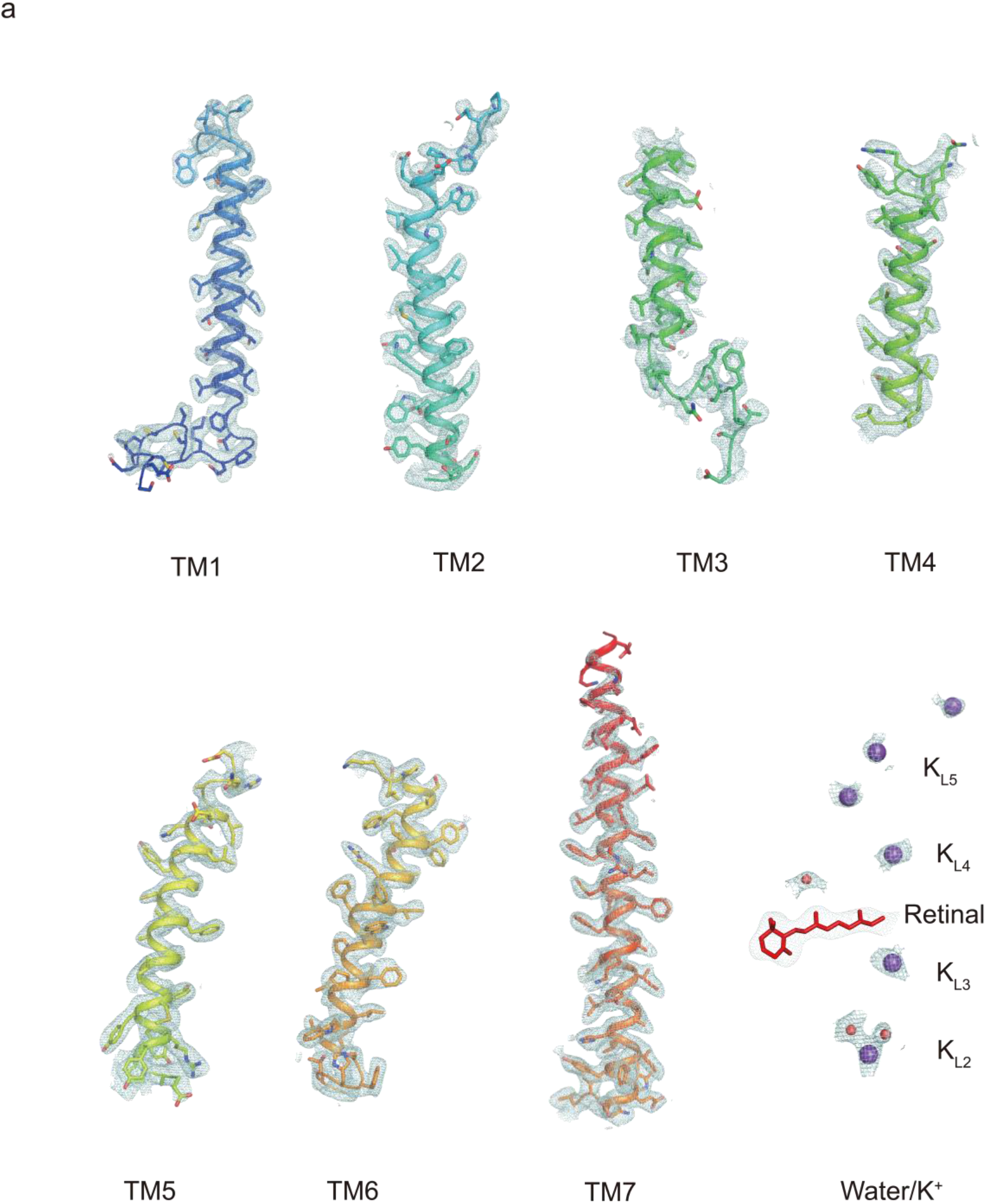
Cryo-EM density of *Bl*CHR2 in potassium environment. **a**, The cryo-EM density of the residues, ion/waters, lipids and the endogenous retinal of *Bl*CHR2 in potassium environment.

**Extended Data Fig. 14.**
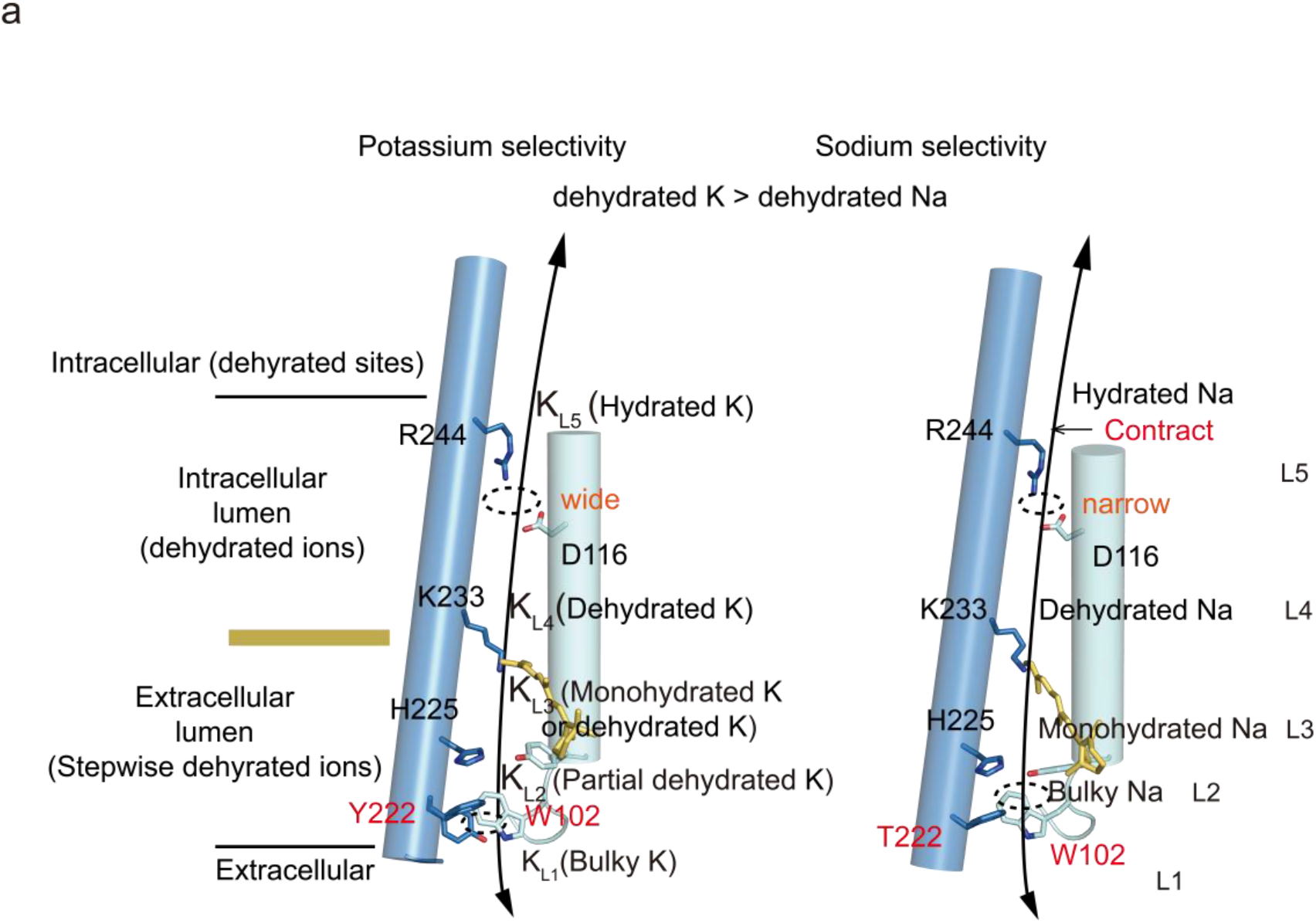
Proposed model for ion selectivity of *Hc*KCR1, *Hc*CCR and *Bl*CHR2. **a**, The stepwise dehydration process in the extracellular side of *Hc*KCR1, *Hc*CCR and *B1*ChR2 play the dominant role in choosing the potassium ions that are more easily dehydrated and the intracellular side of *Hc*KCR1, *Hc*CCR and *B1*ChR2 play the vital role in choosing the right size of ions that are well occupied. The stronger dehydration capacity, the higher potassium selectivity.

**Table S1.** Statistics for data collection and structural refinement.

|  | #1 HcKCR1 | #2 HcKCR1-C110T | #3 HcKCR1-D116N | #4 HcCCR | #5 BICHR2-1 | #6 BICHR2-2 |
| --- | --- | --- | --- | --- | --- | --- |
|  | PDB-8H23 | PDB-8J93 | PDB-8J8U | PDB-8IFU | PDB-8JXO | PDB-8JXP |
|  | EMD-34436 | EMD-36086 | EMD-36080 | EMD-35423 | EMD-36707 | EMD-36708 |
| Data collection and processing |  |  |  |  |  |  |
| Microscope | FEI Titan Krios | FEI Titan Krios | FEI Titan Krios | FEI Titan Krios | FEI Titan Krios | FEI Titan Krios |
| Magnification | 350000 | 270000 | 270000 | 230000 | 350000 | 350000 |
| Voltage (KV) | 300 | 300 | 300 | 300 | 300 | 301 |
| Detector | Falcon 4i | Falcon 4i | Falcon 4i | Falcon 4i | Falcon 4i | Falcon 5i |
| Electron exposure (e-/Å<sup>2</sup>) | 50 | 50 | 50 | 50 | 50 | 50 |
| Defocus range (μm) | -0.9 to -1.2 | -0.9 to -1.2 | -0.9 to -1.2 | -0.9 to -1.2 | -0.9 to -1.2 | -0.9 to -1.3 |
| Pixel size (Å) | 0.35 | 0.45 | 0.45 | 0.57 | 0.35 | 0.35 |
| Symmetry imposed | C3 | C3 | C3 | C3 | C3 | C3 |
| Initial particle images (no.) | 1661K | 1038K | 561K | 406K | 174K | 174K |
| Final particle images (no.) | 207K | 166K | 74K | 266K | 47K | 40K |
| Map resolution (Å) | 2.61 | 2.55 | 2.59 | 2.37 | 2.71 | 2.72 |
| FSC threshold | 0.143 | 0.143 | 0.143 | 0.143 | 0.143 | 0.143 |
| Refinement |  |  |  |  |  |  |
| Initial model | 8H23 | 8H23 | 8H23 | 8H23 | 8H23 | 8H23 |
| Model resolution (Å) | 2.6 | 2.5 | 2.6 | 2.4 | 2.7 | 2.7 |
| FSC threshold | 0.143 | 0.143 | 0.143 | 0.143 | 0.143 | 0.143 |
| Map sharpening B factor (Å<sup>2</sup>) | 95.4 | 84.8 | 68.3 | 87.9 | 86.2 | 81.2 |

**Table S1. Statistics for data collection and structural refinement**
| Model composition |  |  |  |  |  |  |
| --- | --- | --- | --- | --- | --- | --- |
| Non-hydrogen atoms | 6474 | 6474 | 6474 | 6390 | 6294 | 6294 |
| Protein residues | 777 | 777 | 777 | 777 | 777 | 777 |
| Water | 36 | 36 | 36 | 24 | 9 | 9 |
| Ions | 24 | 24 | 24 | 21 | 18 | 18 |
| lipids | 6 | 6 | 6 | 6 | 6 | 6 |
| RET | 3 | 3 | 3 | 3 | 3 | 3 |
| B factors |  |  |  |  |  |  |
| Protein | 33.36 | 29.36 | 33.18 | 31.52 | 48.42 | 44.18 |
| Water | 31.78 | 27.14 | 32.61 | 30.40 | 40.37 | 35.51 |
| R.m.s. deviations |  |  |  |  |  |  |
| Bond lengths (Å) | 0.003 | 0.004 | 0.004 | 0.005 | 0.003 | 0.003 |
| Bond angles (°) | 0.584 | 0.648 | 0.618 | 0.605 | 0.583 | 0.530 |
| Validation |  |  |  |  |  |  |
| Mol Probity score | 1.68 | 1.74 | 1.48 | 1.64 | 1.62 | 1.51 |
| Clash score | 6.17 | 10.04 | 6.33 | 5.58 | 7.33 | 5.56 |
| Poor rotamers (%) | 0.00 | 0.00 | 0.00 | 0.00 | 0.00 | 0.00 |
| Ramachandran plot |  |  |  |  |  |  |
| Outliers (%) | 0.00 | 0.00 | 0.00 | 0.00 | 0.38 | 0.38 |
| Allowed (%) | 1.43 | 1.56 | 1.95 | 1.30 | 2.95 | 2.95 |
| Favored (%) | 98.57 | 98.44 | 98.05 | 98.70 | 96.67 | 96.67 |

## Acknowledgments

We would like to thank the Cryo-EM Facility and High-Performance Computing (HPC) Center of Westlake University for providing cryo-EM and computation support. This work was supported by Westlake Laboratory (Westlake Laboratory of Life Sciences and Biomedicine) and an Institutional Startup Grant from the Westlake Education Foundation to D.P. We also would like to thank all the Cell fate control lab members for their support.

## Author contributions

M.Z. conceived the project. Y.S. and M.Z. designed the experiments. Y.S, L.Z, X.L and T.P prepared the constructs and cell culture. Y.S. prepared the cryo-EM sample and performed the electrophysiological study. Y.S. and M.Z. collected cryo-EM data. M.Z. performed image processing, built the model, analyzed data, and wrote the manuscript draft. D.P. and M.Z. supervised the project. All authors contributed to the manuscript preparation.

## Reporting summary

Further information on research design is available in the Nature Research Reporting Summary linked to this paper.

## Data availability

The coordinate files and cryo-EM maps of *Hc*KCR1, *Hc*KCR1-D116N, *Hc*KCR1-C110T, *Hc*CCR, *Bl*CHR2 class one and *Bl*CHR2 class two have been deposited at Protein Data Bank (Electron Microscopy Data Bank (EMDB)) under accession codes PDB-8H23 (EMD-34436), PDB-8J93 (EMD-36086), PDB-8J8U (EMD-36080), PDB-8IFU (EMD-35423), PDB-8JXO (EMD-36707), PDB-8JXP (EMD-36708) respectively.

